# Charge distribution of coating brush drives inter-chromosome attraction

**DOI:** 10.1101/2024.12.01.626236

**Authors:** Valerio Sorichetti, Paul Robin, Ivan Palaia, Alberto Hernandez-Armendariz, Sara Cuylen-Haering, Anđela Šarić

**Author notes:** Cluster of Excellence Physics of Life, TU Dresden, 01062 Dresden, Germany; Max Planck Institute of Molecular Cell Biology and Genetics, Pfotenhauerstrasse 108, 01307 Dresden, Germany.

## Abstract

The condensation of charged polymers is an important driver for the formation of biomolecular condensates. Recent experiments suggest that this mechanism also controls the clustering of eukaryotic chromosomes during the late stages of cell division. In this process, inter-chromosome attraction is driven by the condensation of cytoplasmic RNA and Ki-67, a charged intrinsically disordered protein that coats the chromosomes as a brush. Attraction between chromosomes has been shown to be specifically promoted by a localized charged patch on Ki-67, although the physical mechanism remains unclear. To elucidate this process, we combine here coarse-grained simulations and analytical theory to study the RNA-mediated interaction between charged polymer brushes on the chromosome surfaces. We show that the charged patch on Ki-67 leads to inter-chromosome attraction via RNA bridging between the two brushes, whereby the RNA preferentially interacts with the charged patches, leading to stable, long-range forces. By contrast, if the brush is uniformly charged, bridging is basically absent due to complete adsorption of RNA onto the brush. Moreover, the RNA dynamics becomes caged in presence of the charged patch, while remaining diffusive with uniform charge. Our work sheds light on the physical origin of chromosome clustering, while also suggesting a general mechanism for cells to tune work production by biomolecular condensates *via* different charge distributions.

## I. INTRODUCTION

When oppositely charged polyelectrolytes are mixed in solution under certain conditions, they undergo phase separation driven by opposite-charge attraction, forming a dense phase: this process is known as *complex coacervation* [1–4]. In recent years, it has been proposed that complex coacervation might be one of the main mechanisms underlying the formation of biomolecular condensates [5–8]. These are dense compartments not bound by membranes found in living cells, like nucleoli, stress granules and P-bodies. Despite the great interest they have generated both in the soft matter and in the biology communities [5, 6, 8–20] the mechanisms of formation of intracellular condensates are still extensively discussed. In particular, a point of contention is whether the interactions driving their formation are mainly unspecific (like charge-charge interactions) or rather specific, *i*.*e*., depending on specific three-dimensional conformation and binding sites [21]. It has also been proposed that these condensed phases, in virtue of their interfacial tension, can generate capillary forces and thus perform work on cellular structures [22–29]. Often, the formation of these condensates involves both RNA and intrinsically disordered proteins (proteins that lack a well-defined three-dimensional structure), which are both ubiquitous in the cellular environment [5, 7, 16, 18, 30–32].

Recent experiments have shown that a process similar to complex coacervation might underlie the phenomenon of chromosome clustering in eukaryotic cell division [33, 34]. A fundamental role in this clustering mechanism is played by the intrinsically disordered protein Ki-67, which localizes on the surface of the chromosomes at the start of cell division [35–38], forming a coat similar to a polymer brush [39, 40]. In the early stages of cell division, Ki-67 is phosphorylated and thus the brush is thought to be negatively charged [34]. This charge, along with potential steric effects, enables Ki-67 to act as a “chromosome surfactant”, ensuring that the chromosomes are individualized (*i*.*e*., that they do not stick to each other) and can properly attach to the mitotic spindle [38]. Towards the end of cell division, however, it is necessary that the chromosomes cluster together to avoid the trapping of large molecules inside the reforming nuclear envelope [33]. In this phase, Ki-67 is dephosphorylated, becoming thus positively charged [34]. This “charge switch” leads to the recruitment of the negatively charged premature ribosomal RNA (pre-rRNA) molecules that are abundant in the cell. These RNA molecules, having an electrical charge opposite to the one of the brush, can stick to the brushes belonging to different chromosomes and thus generate an inter-chromosome attraction [34]. This can be seen as a complex coacervation in which one of the two polymer species (Ki-67) is not free but bound to the chromosome surface. The legitimacy of this interpretation is reinforced by the experimental observation that some free Ki-67 molecules form condensates with RNA in the cytoplasm and enrich on the chromosomes [34]. Under this interpretation, the generation of inter-chromosome attraction bears similarities to the generation of capillary forces induced by the formation of biomolecular condensates [24, 25, 29].

A peculiar characteristic of this particular process is the presence of a highly positively charged patch (CP) on the Ki-67 molecule. When the CP is removed, clustering is largely inhibited [34]; moreover, this highly charged region is evolutionarily conserved in Ki-67 homologues in different species [34]. These observations suggest that there might be a general physical mechanism for which localizing charge in a specific region might lead to a stronger interaction between oppositely charged polymers. This hypothesis is also corroborated by experiments [6, 32, 41–44] and simulations [32, 41, 42, 45] showing that heterogeneous charge distributions can favor the formation of condensates. The presence of the CP is also reminiscent of the “sticker-and-spacer” model of associative polymers [46–48], which has been employed to understand theoretically the formation of biomolecular condensates [16]. According to this model, the condensation of intrinsically disordered proteins and RNA is mainly driven by regions (the “stickers”) which strongly interact with each other, and are separated along the molecule by weakly interacting (“spacer”) regions [16].

From a physical standpoint, the study of the interaction between two charged polymer brushes mediated by an oppositely charged polymer is connected to several phenomena in soft matter. These include counterion-mediated attraction between like-charged surfaces [49] and the interaction between surfaces with adsorbed polymers in good [50–53] and poor [54] solvent conditions. For these systems, attraction is only possible in the presence of poor solvent [54] or if the surfaces are unsaturated [52, 53]. Additionally, polymer bridging between two adsorbing surfaces [55], between charged particles [56], and between a grafted and a non-grafted surface [57] has been investigated. However, it is unknown whether “patchy” charge regions on the grafted polymers, similar to the Ki-67 CP, lead to stronger attraction if the total length and charge of the grafted polymers is left unchanged. Coarse-grained simulations are the ideal tool to address these questions, as they allow to easily and independently tune the physical properties of each molecular species.

In this work, we investigate the mechanism underlying the Ki-67- and RNA-mediated attractive inter-chromosome interactions observed experimentally in eukaryotic cell division, using coarse-grained molecular dynamics simulations. We find that the origin of the attractive interaction is the formation of RNA bridges between the Ki-67 brushes. In particular, we address the question of whether focusing part of the brush polymer’s charge on a highly charged patch leads to stronger inter-chromosome attraction. Importantly, we find that when the brush is uniformly charged, the RNA polymer is adsorbed onto the brushes, making bridging energetically unfavorable unless the two brushes are brought into contact. In the presence of the CP, the RNA localizes at the extremity of the brushes to maximize the interaction with the CPs, but does not fully penetrate into the brush. As a result, the formation of bridges between the two brushes is more likely and we observe a strong attraction between the chromosomes even when the brushes are not in contact.

## II. COARSE-GRAINED MODEL

To model the interaction between mitotic chromosomes mediated by the Ki-67 brush and RNA in late mitosis (schematically represented in Fig. 1A), we adopt a minimal coarse-grained model [58]. The height of the Ki-67 brush which is localized on the chromosome surface is ≈ 90 nm in early mitosis and ≈ 30 nm [34] in late mitosis, and thus much smaller than the diameter of the chromosome arm, which is ≈ 700 nm [59]. Thus, for the purpose of quantifying the local interaction between the chromosome surfaces, we neglect the curvature of the chromosome arms, and represent them as locally flat surfaces. As chromatin is negatively charged, we give to these surfaces a weak uniform negative charge. The Ki-67 brush is represented as flexible polymers grafted on the chromosome surface at regular intervals, whereas RNA molecules are represented as freely diffusing flexible polymers (see Fig. 1B-C for a schematic representation of simulation model). All polymers are represented as chains of spherical beads of diameter *σ* and mass *m* connected by springs. As for the grafting density *ρ*_graft_ of Ki-67, we consider both a sparse brush, with *ρ*_graft_ = 2.0 × 10^−2^*σ*^−2^, and a dense one, with *ρ*_graft_ = 6.3 × 10^−2^*σ*^−2^. The mapping between the simulation units and the experimentally measured values, described in detail in Sec. S1 of the Supporting Information, is chosen in such a way that the bead diameter *σ* (unit of length) corresponds to 11 nm in physical units. Thus, the grafting densities of the sparse brush corresponds to ≈ 170 molecules μm^−2^, and the one of the dense brush to ≈ 520 molecules μm^−2^. These values are comparable with the experimentally measured ones [38].

**FIG. 1.**
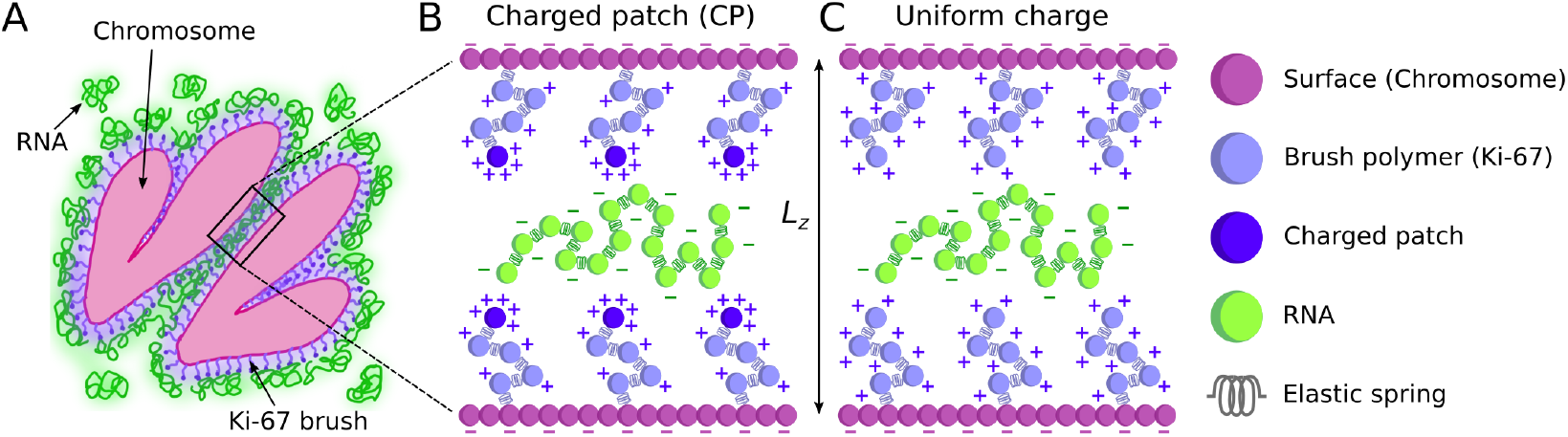
**(A)** Schematic representation of two chromosomes during late mitosis interacting *via* a Ki-67 brush (purple) and RNA (green) **(B-C)** Schematic representation of the simulation model. The chromosomes are modeled as flat charged surfaces kept fixed at a distance *L*_*z*_ from each other. Ki-67 is modeled as 24-bead positively charged polymers bound to the chromosome surfaces. We consider both Ki-67 molecules with a highly charged patch **(B)** and uniformly charged molecules **(C)**, with both types having the same total charge. RNA is modeled as free, 100-bead, highly negatively charged polymers.

The brush polymers are positively charged, mimicking the dephosphorylated state observed in late mitosis. RNA is instead highly negatively charged. Unbonded polymer beads interact via a combination of a short-range repulsive term, mimicking excluded volume interactions [60], and a screened electrostatic potential, with screening length equal to *σ/*2. For the brush polymers, we consider two charge configurations: in one configuration, the farthest bead from the chromosome surface represents the CP and has a charge ≈ 6.5 times higher than that of the other beads, as in the experimental system [34] (Fig. 1B). In the other configuration, the charge of the CP is spread over the whole molecule, resulting in a uniformly charged polymer (Fig. 1C). We stress that the *total* charge of the grafted polymer is the same in both cases. As the experimental CP region’s length is ≈ 1*/*24 of the total length of the brush polymer, the brush polymers are represented for simplicity as 24-bead polymers. The size of the most abundant pre-rRNA molecules differs between different organisms. In humans, pre-rRNA is a mixture of various precursors with different sizes [61, 62]. We take as a reference value the mass of the 45S precursor, ≈ 4.6 MDa (≈ 13000 nucleotides) [63], so that the RNA molecules can be considered to be much larger than the Ki-67 brush polymers (≈ 360 kDa, 3256 amino-acids [64]). Without aiming for a precise mapping of the Ki-67 and RNA length ratio between simulation and experiments, we choose for simplicity to represent RNA as a 100-bead, highly negatively charged polymer. The radius of gyration of an isolated RNA molecule is *R*_*g*,RNA_ = 9.5 ± 0.2. The volume fraction of RNA beads is kept constant and equal to 1%, so that the number of RNA molecules increases when the inter-surface separation *L*_*z*_ is increased.

The solvent is simulated implicitly by means of a Langevin thermostat, and periodic boundary conditions are applied in the horizontal (*x* and *y*) directions. In what follows, we denote with 𝒬 and *τ* respectively the simulation units of charge and time. The bead mass *m* is taken as the unit of mass. The complete details on the simulation model, including the interaction potentials used and the simulation protocol, can be found in Sec. S1 of the Supporting Information.

## III. RESULTS

### A. Coacervation of Ki-67 and RNA generates inter-chromosome attraction

In Fig. 2A-D, we report four representative simulation snapshots, for the sparse (A,B; *L*_*z*_ = 20*σ*) and dense (C,D; *L*_*z*_ = 29*σ*) brush case. To highlight the distribution of the RNA (green), the chromosome surfaces are not shown and the brush polymers (light purple, CP in dark purple) are shown as semi-transparent, except for the CP when present. One can see that when the brush is uniformly charged (Fig. 2A,C), most of the RNA molecules adsorb onto the brushes, with only a small fraction of them occasionally connecting the two surfaces. The spatial organization of RNA changes dramatically in presence of the CP (Fig. 2B,D): here, the RNA is localized in the region between the two brushes, with many RNA molecules forming bridges between these. As we will show below, these bridges lead to inter-surface attraction. Thus, in presence of the CP, the negatively charged RNA acts as a layer of “glue” between the two positively charged brushes, as observed in experiments [34]. Understanding the physical origin of this behavior, and the reason for the striking difference between the uniformly charged brush and the one with the CP, is the main goal of this work.

**FIG. 2.**
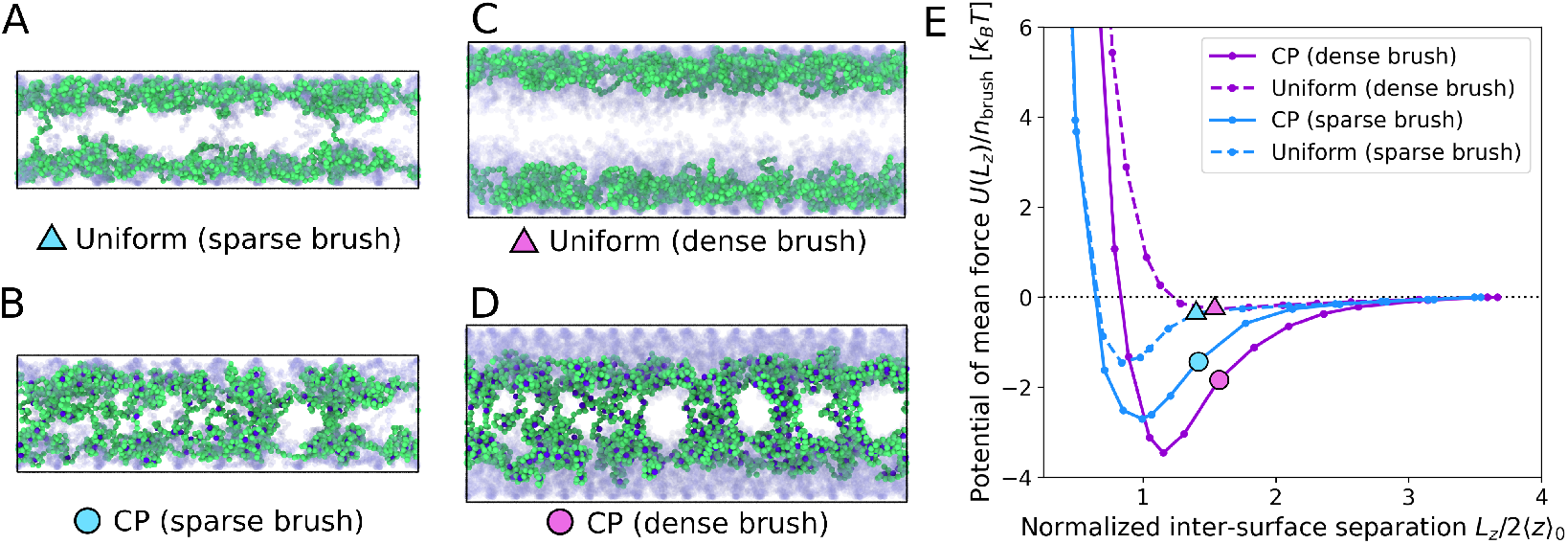
Condensation of Ki-67 and RNA generates inter-chromosome attraction. **(A-D)** Snapshots of the simulated system for sparse (**A**,**B**; *L*_*z*_ = 20*σ*) and dense (**C**,**D**; *L*_*z*_ = 29*σ*) brush. The grafted Ki-67 polymers (light purple) are shown as semi-transparent for clarity. The CP is shown in dark purple, and the RNA polymers in green. **(E)** Potential of mean force [PMF, Eq. (1)] between the grafted surfaces. Simulation results for dense and sparse brushes, with uniform charge and with the CP. The PMF is normalized by the number of grafted polymers per brush *n*_brush_, and the inter-surface separation *L*_*z*_ is normalized by twice the average height of the unperturbed brush ⟨*z*⟩_0_.

To quantify the force acting between the chromosome surfaces, we begin by measuring the potential of mean force (PMF) *U* (*L*_*z*_) acting between them. To obtain this quantity, we measure the *zz* component of the pressure tensor [65], *P*_*zz*_(*L*_*z*_), and obtain from this the force *F*_*z*_(*L*_*z*_) = *AP*_*zz*_(*L*_*z*_) acting between the two chromosome surfaces (*A* being the surface area). To make sure that the system is as close as possible to a steady state, we only consider the average value of *P*_*zz*_ measured at long times. The sign of *F*_*z*_ determines whether the force acting between the surfaces is attractive (*F*_*z*_ *<* 0) or repulsive (*F*_*z*_ *>* 0). The PMF is then defined as

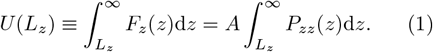

Note that, based on this definition, the PMF is zero for infinite inter-surface separation.

In Fig. 2E we show the PMF as a function of the inter-surface separation *L*_*z*_. The latter quantity is normalized by 2 ⟨*z*⟩_0_, with ⟨*z*⟩_0_ the mean height of the unperturbed (RNA-free) brush (see Sec. S3 A of the Supporting Information). Moreover, to facilitate the comparison between the sparse and dense brush systems, we divide *P*_*zz*_ by the number of brush polymers per surface, *n*_brush_. One can see that the PMF for the sparse brush displays an attractive well [*U* (*L*_*z*_) *<* 0] both when uniformly charged and with the CP. However, the attractive well is deeper with the CP: thus, the CP leads to stronger attraction between the chromosome surfaces, in agreement with experiments with similar brush densities [34]. The most striking difference between uniform charge and CP, however, is observed for the dense brush: with the CP, *U* (*L*_*z*_) displays an even deeper well, indicating strong attraction. For the uniformly charged brush, instead, the attractive well disappears almost completely due to the RNA polymers adsorbing onto the brushes (Fig. 2A,C).

Another notable difference between the uniformly charged brush and the one with the CP is that, if the grafting density is increased in the CP system (solid lines), the depth of the attractive well increases, *i*.*e*. the attraction become stronger. For the uniformly charged brush (dashed lines), however, the opposite is true: when the grafting density is increased, the attractive well almost disappears. Intuitively, this is due to the fact that, for higher grafting density, even more RNA polymers are adsorbed onto the brushes and do not form bridges.

To estimate what would be the equivalent strength of the inter-chromosome attraction in physical units, we compute the maximum attraction force between mitotic chromosomes. For the sparse brush, which has a Ki-67 density which is closer to the one of the experimental system, the magnitude of the force exerted by each grafted polymer close to the minimum of the PMF is |*F*_*z*_| */n*_brush_ ≈ 0.3*k*_*B*_*T/σ* (See Sec. S3 B of the Supporting Information, in particular Fig. S4). Assuming *k*_*B*_*T* = 310 K and *σ* ≈ 11 nm (see Sec. S1 of the Supporting Information), we have |*F*_*z*_| */n*_brush_ ≈ 0.1 pN. The number of Ki-67 molecules on the chromosome surface can be estimated by assuming a surface density *ρ*_graft_ ≈ 210 μm^−2^ [38]. Assuming a contact area of the order of 1 μm^2^ for two interacting chromosomes, we have thus *n*_brush_ ≈ 200 molecules, so that the total force acting between the chromosomes would be |*F*_*z*_| ≈ 20 pN in physical units (corresponding to a pressure magnitude |*P*_*zz*_| ≈ 20 Pa). A larger contact area would lead to a larger force; nevertheless, this estimate is comparable the typical forces exerted on mitotic chromosomes by the spindle, which are of the order of tens to hundreds of pN [66–68].

The result that the CP leads to stronger and more long-ranged attraction compared to the uniform charge remains true also upon changing the RNA length and increasing its concentration (see Sec. S3 C and S3 D of the Supporting Information). In general, increasing the RNA concentration leads to the adsorption of more RNA polymers onto each brush, which reduces the electrostatic repulsion between the brushes. However, in the CP case, the presence of more RNA molecules also leads to saturation of the CPs, and thus to a decrease of the attraction. These effects are discussed in detail in Sec. Sec. S3 C of the Supporting Information.

### B. Inter-chromosome attraction is mediated by RNA bridges

In Sec III A, we have shown that there is an attractive potential of mean force (PMF) between the two surfaces, and that the strength and range of the attraction are significantly larger in the CP case than in the uniform charge case. Below, we show that this attractive interaction is generated by the formation of RNA bridges between the two brushes, and that these bridges are more likely to form in presence of the CP. In order to do so, we start by comparing the time evolution of the *zz* component of the pressure tensor, *P*_*zz*_, to the time evolution of the fraction of bridging RNAs. We only discuss here the case of the dense brush, which displays the most striking difference between uniform charge and CP. A similar analysis for the sparse brush, holding the same qualitative results, is reported in Sec. S3 E of the Supporting Information.

In Fig. 3A, we show the time evolution of *P*_*zz*_ for the CP case, for four different values of the inter-surface separation *L*_*z*_. We normalize *P*_*zz*_ by the number of Ki-67 polymers per brush, *n*_brush_ as done in Fig. 2E for the PMF. At short times, the value of *P*_*zz*_ is negative, implying attractive interaction between the two surfaces. For *L*_*z*_ *>* 24*σ*, the value of *P*_*zz*_ gradually increases over time, signifying a reduction of the attractive interaction. For the largest separation considered, *P*_*zz*_ quickly relaxes to zero (no net force between the surfaces). For the smallest separation, on the other hand, *P*_*zz*_ remains at a constant negative value, implying that there is stable attraction between the surfaces.

**FIG. 3.**
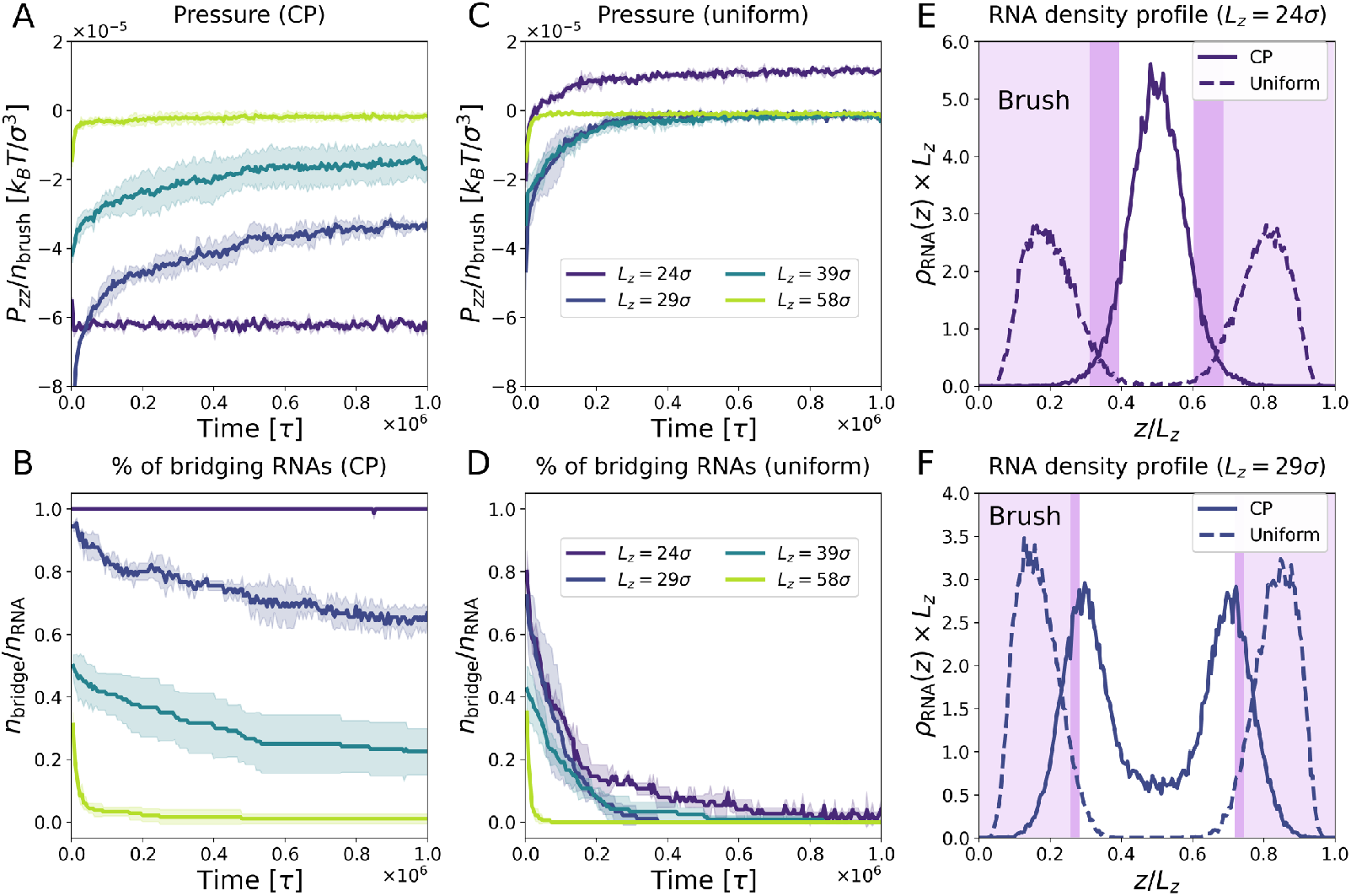
Inter-surface attraction is mediated by RNA bridges. **(A-C)** Time evolution of the *zz* component of the pressure tensor *P*_*zz*_, normalized by the number of grafted polymers per brush *n*_brush_. **(A)** CP, dense. **(C)** Uniform, dense. **(B**,**D)** Time evolution of the fraction of bridging RNAs, *n*_bridge_*/n*_RNA_. **(B)** CP, dense. **(D)** Uniform, dense. **(E-F)** Normalized *z*-density profile of RNA for the dense brush with inter-surface separation *L*_*z*_ = 24*σ* **(E)** and *L*_*z*_ = 29*σ* **(F)**. The shaded area represents the extension of the brush, with the lighter shade for the uniformly charged brush and the darker shade for the CP one.

To show that this attractive interaction is due to the formation of RNA bridges between the two brushes, we study the time evolution of the fraction of bridging RNAs, *n*_bridge_*/n*_RNA_. We define a bridge as any RNA polymer whose end monomers are neither both in the portion 0 *< z < L*_*z*_*/*2 of the box nor in *L*_*z*_*/*2 *< z < L*_*z*_. This definition is well suited to our system, where each RNA molecule is usually either bridging or adsorbed onto the brush, and in our simulations it turned out to be the one that most reliably captures the correct number of bridges. We also considered an alternative definition which explicitly takes into account the brush extension, finding very similar results (see Sec. S3 F of the Supporting Information).

In Fig. 3B, we show the fraction of bridging RNAs as a function of time for the same systems as in Fig. 3A. One can see that there is a very clear correlation between the number of bridges formed between the surfaces and the value of the pressure: when almost no bridge is present, *P*_*zz*_ = 0 and there is no inter-surface attraction. Instead, when many bridges are formed, *P*_*zz*_ *<* 0 and attraction is present. For *L*_*z*_ = 24*σ*, for which we had observed a stable negative value of *P*_*zz*_, almost all the RNAs are bridging. Thus, we conclude that RNA-mediate bridging is indeed at the origin of the attractive force between the surfaces.

In Fig. 3C-D, we present the same analysis for the uniform charge case. For *L*_*z*_ *>* 24*σ*, the pressure rapidly grows to zero, while at the same time the fraction of bridging RNAs also decays to zero. We thus conclude that for the uniformly charged brush, not only much fewer bridges are formed initially, but their lifetime is significantly shorter. For *L*_*z*_ = 24*σ, P*_*zz*_ grows to a positive value, implying that there is a inter-surface *repulsion*, in agreement with the PMF analysis (Fig. 2). We note that, after an initial fast decrease, the number of bridges decays approximately exponentially for the uniform brush, whereas the decay is slower than exponential in the CP case (see also Sec. S3 G of the Supporting Information). We attribute this non-exponential relaxation to the fact that each RNA bridge has to break bonds with multiple CPs to be released [69].

The dramatic difference between the CP and the uniform case is due to the different spatial arrangement of the RNA polymers in the two cases. This is shown quantitatively in Fig. 3E and F, where we report the normalized density profile of RNA along the *z* axis, *ρ*_RNA_(*z*), for *L*_*z*_ = 24*σ* and *L*_*z*_ = 29*σ* respectively. The shaded areas represent the extension of the brush. One can see that in the uniform case, RNA is mostly adsorbed onto the brush, whereas in the CP case there is a high concentration of RNA (bridging) in the zone between the two brushes. For the detailed evolution of *ρ*_RNA_(*z*) as a function of *L*_*z*_, and the density profiles of the brush polymers and of the CPs, see Sec. S3 H of the Supporting Information.

### C. A localized charged patch is required for stable bridging

To conclude our analysis of the bridging mechanism, we show with simulation data and theoretical arguments that a localized CP is a required condition for bridging. Without it, RNA completely adsorbs onto the brush, which prevents bridging. To this end, we study how the fraction of bridging RNA polymers depends on the inter-surface separation *L*_*z*_. As for the PMF, we take the average value of *n*_bridge_*/n*_RNA_ measured at long times. In Fig. 4A, we report this quantity for all the studied systems, including the ones with sparse brush. We observe that in the uniformly charged brush case (dashed lines), the fraction of bridging RNAs quickly drops to zero as soon as the brushes are not in contact. One can see this by comparing the curve to the shaded areas, which represent twice the extension of the unperturbed brush, 2 ⟨*z*⟩_0_, in the uniform case. Thus, for the uniformly charged brush, bridging is only possible if the two brushes are in contact: as soon as they are separated, the fraction of bridging RNAs drops to zero. The CP case is dramatically different (solid lines): here, the fraction of bridging RNAs is significantly different from zero even at distances larger than the contact distance between the brushes. Thus, the localized CP allows the formation of long-range bridges that do not require the two brushes to be in contact to exist. Moreover, we also observe a marked difference between the dense and sparse case, which is not observed in the case of uniform charge.

**FIG. 4.**
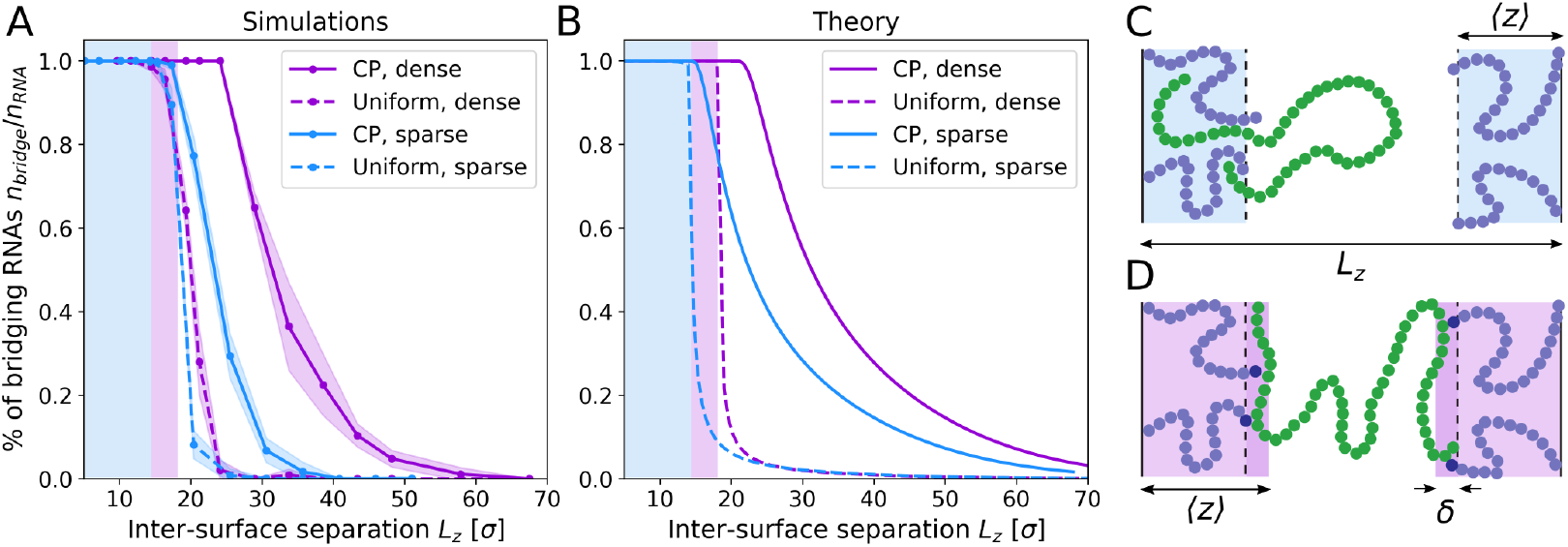
A localized charged patch is required for stable bridging. **(A**,**B)** Fraction of bridging RNAs as a function of the inter-surface separation *L*_*z*_ for different systems. **(A)**: Simulations. **(B)**: Theoretical prediction. Shaded areas in **(A)** and **(B)** represent twice the extension of the unperturbed brush, 2 ⟨*z*⟩ _0_, for the uniform dense (pink) and the uniform sparse (blue) brush. For the uniformly charged brush, the fraction of bridging RNAs suddenly drops to zero as soon as the brushes are not in contact. In contrast, with the CP, the fraction of bridging RNA is significantly different from zero even when the brushes are not in contact. The theoretical prediction show good qualitative agreement with the simulation results. **(C**,**D)** Schematic representation of the system used for the theoretical model for the uniform **(C)** and CP **(D)** case. In the uniform case, the RNA (green) can penetrate into the brush, whereas in the CP case it can only penetrate in a thin slab of thickness *δ* ≈ *σ* where the CPs are localized.

To understand the physical reason why the CP is required for bridging, we develop a simplified mathematical model of the simulated system (detailed in Sec. S2 of the Supporting Information). In this model, the RNA molecules are represented as ideal chains [70] of contour length *𝓁*, confined between two infinite surfaces placed at a distance *L*_*z*_ from each other. Each polymer brush is modeled as a region of thickness ⟨*z*⟩ adjacent to the surface, as shown in Fig. 4C-D. For simplicity, we assume here that ⟨*z*⟩ remains constant when *L*_*z*_ is changed. As discussed in Sec. S3 B of the Supporting Information, this assumption is approximately valid as long as the two brushes do not interpenetrate. We then distinguish two regions within the brush: one of thickness *h*, adjacent to the plates, that RNA is excluded from (see Fig. 3E-F); and a second of thickness *δ* into which RNA can penetrate, and where it experiences an enthalpic gain quantified by the excess free energy − Δ*F <* 0 per bead due to electrostatic interactions with the brush. In the case of the brush with the CP, the RNA polymers do not penetrate significantly into the brush, as shown in Fig. 2B,D and quantified in Figs 3E-F: thus, only the region of space between the brushes is assumed to be available to the RNA molecules and *h* ≈ ⟨*z*⟩. Electrostatic interactions are localized around the CP, so we assume *δ ≈ σ*. In the case of the uniformly charged brush, on the other hand, the RNA molecules penetrate into the brush, with which it continuously interacts, and thus *h ≈* 0 and *δ ≈*⟨*z*⟩.

The main result of this model is an analytical expression for the equilibrium fraction of bridging RNA molecules as a function of the inter-surface separation:

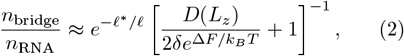

where and *D* is the *z*-width of the region of space available to the RNA, so that *D* = *L*_*z*_ for the uniform charge case and *D* = *L*_*z*_ *™* 2 ⟨*z*⟩ for the CP case. Eq. (2) was obtained under the assumption that the RNA monomers bind strongly to the brush, 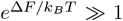. The length *ℓ*^*^ represents the critical RNA length that, for a given distance *D*, separates the regime in which no bridges are formed (*ℓ < ℓ*^*^) from the one where most RNA molecules are bridging (*ℓ > ℓ*^*^). The value of *ℓ*^*^ depends in general on *D, h* and Δ*F*, but in the regime relevant for attraction one finds 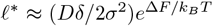. By equating *ℓ*^*^ and *ℓ* we thus find the critical value of *D* above which most bridges are broken, which corresponds to the range of the attractive interaction between plates:

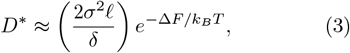

with the corresponding critical inter-surface separation being 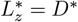 for the uniformly charged brush and 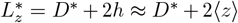 for the CP brush. In the CP case, while the interaction energy between individual RNA beads and CPs is high (≈6 *k*_*B*_*T*), only a small fraction of beads per RNA chain can bind to the brush through the CPs. Employing Flory-Huggins theory [70] (see Sec. S2 A of the Supporting Information), we estimate that the excess free energy per bead is in this case Δ*F≈*3 *k*_*B*_*T*. With these parameters, we find that the system undergoes “partial coacervation”: the RNA is not fully adsorbed onto the brush, and bridging is possible over a distance 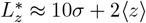 between the brushes.

For the of uniformly charged brush, while electrostatic interactions between individual RNA and brush beads are weaker than in the CP case, adsorption of the RNA polymers onto the brush in entropically favored, resulting in a higher free energy per bead Δ*F*. We find Δ*F ≈* 5 *k*_*B*_*T*, and the system is then in the “complete coacervation” regime: RNA mostly adsorbs onto the brush, and bridging can only occur when brushes are in direct contact. We note that increasing the charge of the brush polymers would not result in a significant increase of *n*_bridge_, as the RNA polymers would have an even stronger tendency to fully condensate with the brush. We show this in Sec. S3 I of the Supporting Information. Additionally, in Sec. S3 J of the Supporting Information we show that our results are robust with respect to changes of the electrostatic interaction strength and screening length.

In Fig. 4B, we report the theoretical predictions for the fraction of bridging RNAs [Eq. (2)]. Despite the approximations of the analytical model, we observe good qualitative agreement between theory and simulations. A detailed discussion of the assumptions of this mathematical model and the details of the calculations can be found in Sec. S2 of the Supporting Information.

### D. The charged patch induces dynamical caging of the RNA molecules

The presence or absence of the CP strongly influences also the dynamics of RNA. In order to quantify this, we measure the mean-squared displacement (MSD) [71] of the RNA molecules, 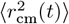, computed by tracking the motion of their center of mass. For a freely diffusive polymer without hydrodynamics, the MSD of RNA is given by the Rouse model [72], which predicts normal diffusion, 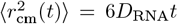, with a diffusion coefficient *D*_RNA_ = *k*_*B*_*Tσ/*(*ζN*_RNA_). Here *ζ* is the viscous friction each monomer experiences and *N*_RNA_ the number of beads in each RNA polymer. Using the numerical values of the parameters, we predict *D*_RNA_ = 10^−3^*σ*^2^*/τ* for a freely diffusing RNA molecule. We start the measuring of the MSD at the time *t*_0_ = 5 × 10^5^*τ* to minimize the effect of the initial transient in which the RNA bridges are formed and then progressively broken.

In Fig. 5A, we show the MSD of the RNAs for the sparse brush system, for three different values of the inter-surface separation *L*_*z*_. For the uniformly charged brush (dashed lines), the MSD is independent of *L*_*z*_ Initially, the RNA polymers follow the Rouse model prediction, with *ζ* = 10*m/τ* the friction of the solvent. In this early times regime, the dynamics is very similar to that of free RNA polymers in the absence of the brushes (free RNA, solid gray line). At later times, however, the RNA molecules start to interact more strongly with the brush. This results in an increase of the effective viscous friction, which becomes approximately ten times larger *ζ*_eff_ ≈ 10*ζ*. The dynamics remains however diffusive.

**FIG. 5.**
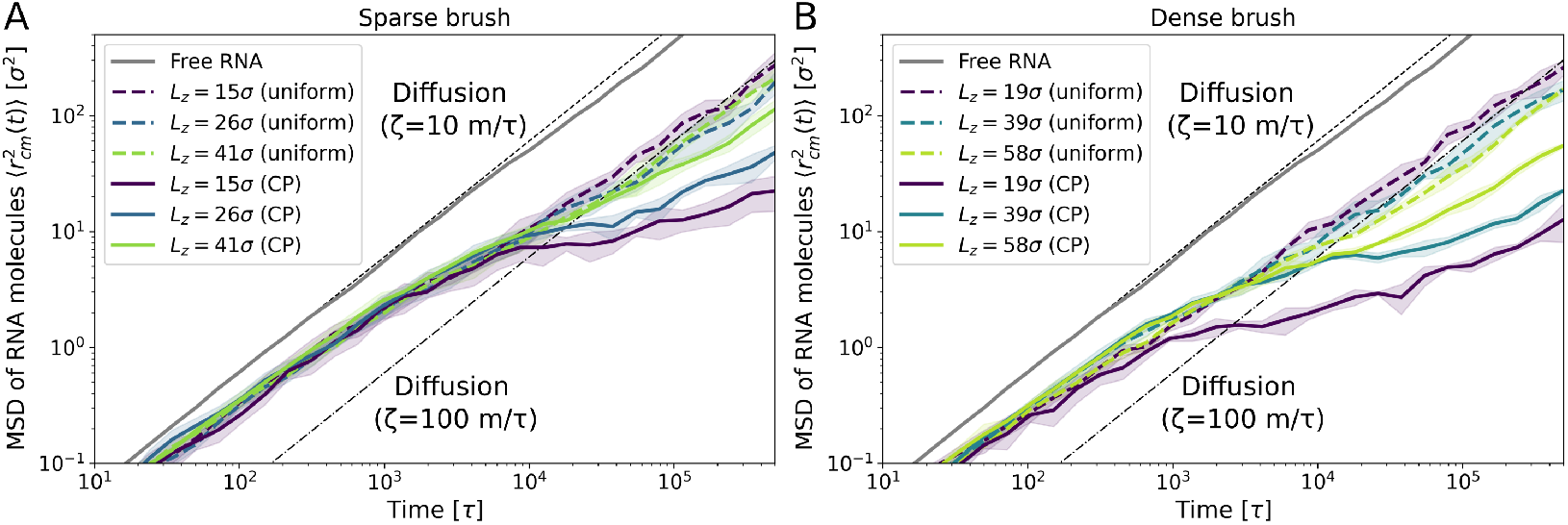
The charged patch induces dynamical caging of the RNA molecules. MSD of the RNA molecules for sparse **(A)** and dense **(B)** brush, for different values of the inter-surface separation *L*_*z*_. Dashed line: Rouse model prediction for a freely diffusive polymer in the absence of hydrodynamic interactions, 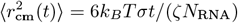. Here *σ* is the monomer diameter, *ζ* = 10*m/τ* is the viscous friction and *N*_RNA_ = 100 is the number of beads of an RNA molecule. Dash-dotted line: Rouse model prediction with effective viscous friction *ζ*_eff_ = 10*ζ* = 100*m/τ*. The solid gray line is the MSD of free RNA polymers at the same density as the one used in our system.

The long-time dynamics of RNA changes significantly in the presence of the CP (solid lines), with a marked subdiffusive transient in the MSD (*i*.*e*., 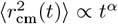, with *α <* 1) signaling the presence of strong caging of the RNA molecules. This behavior (sometimes referred to as “sticky Rouse” dynamics) is due to the fact that an RNA polymer needs to break the many strong bonds formed with the CPs in order to diffuse, and it is a common feature of systems of polymers with binding sites [47, 48, 73–79]. The degree of caging also increases when *L*_*z*_ is decreased, due to the formation of bridges that interact with even more CPs.

We note that the observed caging effect is not due to entanglements [72], as it is clear from the fact that it is absent for the uniformly charged brush. However, by analyzing the MSD of the RNA monomers, we uncover a subdiffusive regime with exponent numerically close to the one found for entangled polymers [72, 80] (see Sec. S3 K of the Supporting Information). We thus conclude that CP leads to caging of the RNA molecules, and to a marked dynamical slowing down. Our results show how localized CPs can be a determinant factor contributing to the high viscosity of some biomolecular condensates, which have been shown to behave as glassy viscoelastic materials [81–84].

## IV. DISCUSSION AND CONCLUSION

Employing a minimal coarse-grained model of charged polymers, we have studied the interaction between mitotic chromosome surfaces, as mediated by Ki-67 brushes and RNA. In our model, inspired by recent experiments [33, 34], the negatively charged RNA is able to generate inter-chromosome attraction by forming bridges between the positively charged brushes present on the chromosome surfaces. The RNA molecules attached to the brushes pull the chromosome surfaces together and generate between them an attractive force. Importantly, we show that when the brush is uniformly charged, bridging is only possible when the two brushes are in contact, and the bridges are easily broken. On the other hand, in the presence of highly charged patches (CP), bridging becomes possible also when the brushes are not in contact. This leads additionally to more long-lived bridges and to a stronger inter-chromosome attraction. The observation that a “patchy” charge distribution can lead to a stronger interaction between proteins and RNA is in line with recent *in vivo* [34] experiments, and also is also reminiscent of the “sticker-and-spacer” model for the formation of biomolecular condensates [16]. To summarize, both in the CP and uniform case a coacervate is formed that acts as a sort of “glue” between the chromosomes. However, in the uniform case this glue is only effective when the two brushes are at contact, whereas the CP makes the glue stronger while also enabling it to act at a longer range.

The maximum attractive force measured from the simulation corresponds to ≈ 20 pN in physical units for a contact area of order 1 µm^2^. This force is comparable to the typical ones exerted by the mitotic spindle (of order of ten to hundreds of pN [66–68]), which suggests that it might act concurrently with spindle forces to drive chromosome clustering during mitotic exit. Our estimate of the maximum attractive force is larger than recent measurements of the forces exerted by biomolecular condensates, which reported values between 0.1 and 1 pN [24, 29]. However, we note that these studies measured the force exerted by condensates on single strands of DNA [24] and chromatin [29]. In our case, the force exerted by each brush polymer is ≈ 0.1 pN, which results in a total force of a ≈ 20 pN for the ≈ 200 polymers covering a 1 µm^2^ area. Moreover, this estimate depends on the assumptions of our model, in particular regarding the charge of pre-rRNA, which has yet to be precisely measured in experiments.

From a more general point of view, the physical mechanism that we described bears strong similarity to the process of complex coacervation, the phase separation of oppositely charged polymers [1, 3, 4], which has been suggested to be one of the main drivers of the formation of biomolecular condensates [5–8]. Importantly, biomolecular condensates are known to be able to exert capillary forces, performing work on the surrounding environment [22–29]. As previously suggested [34], the attraction mechanism here reported is somewhat reminiscent of a capillary interaction resulting from the coacervation of Ki-67 and RNA. Despite the similarity between our bridging mechanism and capillary interactions, however, the scaling of the force with distance is different in the two cases [85, 86] (see Sec. S3 L of the Supporting Information), which shows that these are distinct mechanisms. However, capillary forces might still play a role *in vivo*.

In the case of uniformly charged brush, a “complete coacervation” of Ki-67 and RNA takes place, in that RNA is completely adsorbed onto the brushes. Under these conditions, bridging is thermodynamically unfavorable, and inter-chromosome attraction is low or even zero. In presence of the CP, the RNA undergoes instead a “partial coacervation” with the CPs only, and does not completely adsorb onto the brush, making bridging –and thus attraction– possible. This “partial condensation” is qualitatively reminiscent of the micro-phase separation observed in systems of block copolymers [87–91], although the latter is driven by the thermodynamic incompatibility of different polymer regions.

An additional prediction of our simulations is that the presence of the CP leads to significant dynamic slowing down of the RNA molecules. This is due to the many strong bonds formed between an RNA molecule and the CPs, which must be broken in order for the molecule to diffuse (“sticky Rouse” dynamics [47, 73, 74]). This generates a “caging” effect, resulting in a marked dynamical slowing down similar to the one observed for entangled polymers – despite our system being unentangled. Localized charged patches can thus be one of the mechanism that contribute to the high viscosity of biomolecular condensates measured in experiments [81, 82].

We note that in our minimal coarse-grained model, the effect of ions is encoded implicitly through an electrostatic screening term. This is justified by the observation that electrostatic interactions are strongly screened at physiological ion concentrations [92]. For these typical concentrations, we expect our main result, that the CP leads to stronger and more long-range attraction between the chromosomes, to remain valid. This is also supported by the observation that our results are robust with respect to changes over a range of parameters representing electrostatic interactions (see Supporting Information). However, there might be biological perturbations, such as hypertonic/hypotonic shocks, that might change this picture. Hypertonic shocks, for example, would lead to an increase of the ionic strength of cytoplasm and thus to a weakening of the electrostatic interactions due to increased screening. This would not only weaken the bridging mechanism, but also alter the binding of Ki-67 to chromatin and possibly even chromatin’s compaction state. Conversely, decreasing the ionic strength would make the electrostatic interactions more long-ranged, possibly smearing out the difference between CP and uniform charge. Exploring these effects would require a more detailed simulation model, including long-range electrostatics and explicit ions and counterions. Another interesting question concerns the degree of charge localization required to observe the reported phenomenon, as one might expect the difference between uniform charge and CP to gradually disappear as the CP becomes larger. These questions open interesting avenues for future investigations on the role of charge localization in biomolecular condensates.

In conclusion, we have shed light on the microscopic mechanism of inter-chromosome attraction resulting from the condensation of Ki-67 and RNA in mitotic exit [33, 34], elucidating the physical reasons for which a localized charge distribution leads to stronger and more long-ranged attraction. The mechanism here described suggests a general avenue for the cell to control the forces exerted by condensates using proteins with different charge patterns, tuning the strength and range of the interaction by changing *e*.*g*. the protein phosphorylation state.

## ACKNOWLEDGMENTS

This work has received funding from the European Research Council (ERC) under the European Union’s Horizon 2020 research and innovation program (grant agreement No. 802960) and support by the German Research Foundation (DFG project number 402723784). V. S. and A.Š. acknowledge funding from The Vallee Scolarship. I.P. and P.R. acknowledge funding from the European Union’s Horizon 2020 research and innovation program under the Marie Skłodowska-Curie grant agreement No. 101034413. A.H.-A. received a PhD fellowship from the Boehringer Ingelheim Fonds.

