## Supporting Information for "Charge distribution of coating brush drives inter-chromosome attraction"

### CONTENTS

S1. Simulation model 1
- A. Interaction potentials 1
- B. Charge distribution and simulation protocol 2
- C. Mapping to physical units 3

S2. Analytical model 3
- A. Flory-Huggins approach to coacervation 4
- B. Model definition 5
- C. Bridging probability 5
- D. Charged patch case 7
- E. Scaling 9
- F. Uniformly charged brush 9
- G. Results and discussion 9

S3. Additional simulation data 10
- A. Unperturbed brush height 10
- B. Brush height and force between the two grafted surfaces 10
- C. Effect of increasing RNA concentration 11
- D. Effect of changing RNA length 14
- E. RNA bridging for the sparse brush 14
- F. Alternative definition of bridge 15
- G. Decay of number of bridges in time 15
- H. Density profiles of RNA, brush monomers and CP along the $z$ axis 16
- I. PMF dependence on brush polymer charge (dense uniform case) 17
- J. PMF dependence on the electrostatic interaction strength and screening length 18
- K. Subdiffusive dynamics of RNA monomers 20
- L. Comparison to capillary forces 21

References 22

### S1. SIMULATION MODEL

#### A. Interaction potentials

To model the interaction between mitotic chromosomes mediated by Ki-67 and RNA, we perform $NVT$ (constant number of particles, volume and temperature) molecular dynamics (MD) simulations of a coarse-grained model (Fig. 1 of the main text). Since we focus here on the microscopic origin of the attraction force, we model the chromosomes as two rigid flat surfaces made of spherical beads arranged on a squared lattice. Locally approximating the chromosome with a flat surface is further justified by the fact that the diameter of a mitotic chromosome arm is of the order of 700 nm [1], whereas the height of the dephosphorylated Ki-67 brush is $\approx 90$ nm during early mitosis and $\approx 30$ nm in mitotic exit [2].

The Ki-67 molecules are modeled as bead-spring polymers grafted to the chromosomes surfaces, whereas RNA molecules, also modeled as bead-spring polymers, are not bound and can freely diffuse in the system. The interaction between all the beads (belonging to the chromosome surfaces, to the Ki-67 and to the RNA) combines an excluded volume part, which models steric interactions, to a screened electrostatic part. The excluded volume interaction is modeled using the WCA potential [3],

$$U_{\mathrm{WCA}}(r) = \begin{cases} 4\epsilon\left[(\sigma/r)^{12} - (\sigma/r)^{6}\right] + \epsilon & r \leq 2^{1/6}\sigma \\ 0 & \text{otherwise,} \end{cases} \tag{S1}$$

[*] Current addresses: Cluster of Excellence Physics of Life, TU Dresden, 01062 Dresden, Germany; Max Planck Institute of Molecular Cell Biology and Genetics, Pfotenhauerstrasse 108, 01307 Dresden, Germany.

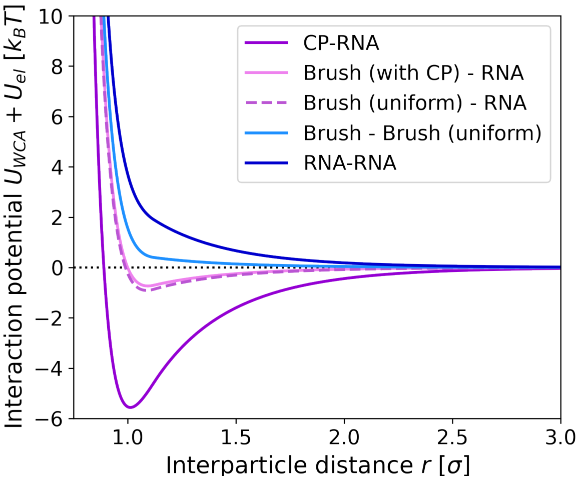


FIG. S1. The total interaction potential $U_{\rm WCA} + U_{\rm el}$, sum of Eq. (S1) (excluded volume) and Eq. (S2) (screened electrostatic), for the different particle types and charge configurations. The charge of an RNA bead is $q_{\rm RNA} = -10\mathcal{Q}$, that of the CP is $q_{\rm CP} = 24.15\mathcal{Q}$. The charge of the Ki-67 (brush) beads is $q_0 = 3.72\mathcal{Q}$ in the CP case and $4.57\mathcal{Q}$ in the uniform case.

with $\sigma$ the bead diameter and $\epsilon$ the interaction energy. Throughout this work, we take as units of energy, length and mass respectively $\epsilon$, $\sigma$ and $m$ (the mass of the beads comprising the Ki-67 and RNA polymers). Additionally, $\epsilon = k_BT$ throughout. All other units are derived from these, so that the unit of time is $\tau = \sqrt{m\sigma^2/\epsilon}$, the one of force is $\epsilon/\sigma$ and the one of pressure is $\epsilon/\sigma^3$. The unit of charge is $\mathcal{Q} = \sqrt{4\pi\varepsilon_0\sigma\epsilon}$, with $\varepsilon_0$ the vacuum permittivity. The screened electrostatic potential is given by

$$U_{\rm el}(r) = \begin{cases} K q_i q_j \exp(-r/\lambda)/r & r \leq 3\sigma \\ 0 & \text{otherwise}, \end{cases} \tag{S2}$$

where $q_i, q_j$ are the charges of the two interacting particles, $K$ is a constant controlling the strength of the electrostatic interaction and $\lambda$ is the screening length. The choice of a screened electrostatic potential is justified by the observation that, for typical intracellular ionic concentrations, electrostatic interactions are heavily screened [4]. Here, we choose $K = 0.2\epsilon\sigma\mathcal{Q}^{-2}$ and $\lambda = \sigma/2$, and apply a cutoff $3\sigma$. Ours being a minimal coarse-grained model, the electrostatic potential (S2) is not an exact description of microscopic electrostatic interactions, and should be taken as a qualitative representation of the latter. The total interaction potential $U_{\rm WCA} + U_{\rm el}$ is shown in Fig. S1 for different types of bead pairs.

Bonded neighbors in the same polymer chain interact only *via* the excluded volume potential Eq. (S1) plus a finite extensible nonlinear elastic (FENE) potential:

$$U_{\rm FENE}(r) = -\frac{K_{\rm FENE} r_0^2}{2} \ln\left[1 - (r/r_0)^2\right], \tag{S3}$$

with strength $K_{\rm FENE} = 30\epsilon\sigma^{-2}$ and equilibrium length $r_0 = 1.5\sigma$. These values are chosen to prevent chain crossing [5]. The equilibrium bond length with these parameters is $\approx 0.97\sigma$.

#### B. Charge distribution and simulation protocol

The surfaces, which are fixed in space at a distance $L_z$ from each other, are made of spherical beads with diameter $\sigma$ and charge $q_{\rm chr} = -0.5\mathcal{Q}$, arranged on a square lattice with surface density $\sigma^{-2}$. We graft to each surface $n_{\rm brush}$ Ki-67 molecules of $N_{\rm brush} = 24$ beads, which are regularly arranged on a square lattice and bound to the surface by FENE bonds [Eq. (S3)]. The distance between two neighboring grafting sites determines the brush grafting density $\rho_{\rm graft}$ and thus the conformation of the grafted polymers [6, 7]. We consider two grafting densities, $\rho_{\rm graft} = 2.04 \times 10^{-2}\sigma^{-2}$ (sparse brush) and $6.25 \times 10^{-2}\sigma^{-2}$ (dense brush), corresponding to separations respectively $7\sigma$ and $4\sigma$ between neighboring grafting sites. The number of Ki-67 molecules per surface is $n_{\text{brush}} = 100$ for the sparse brush and 324 for the dense one.

RNA molecules are modeled as freely diffusing polymers of $N_{\text{RNA}} = 100$ beads. The radius of gyration of an isolated RNA molecule is $R_{g,\text{RNA}} = 9.5 \pm 0.2$. The number density of RNA molecules is kept constant and equal to $\rho_{\text{RNA}}^{\text{mol}} = 2 \times 10^{-4}\sigma^{-3}$, so that the volume fraction of RNA beads is $\phi_{\text{RNA}} = (\pi/6)\rho_{\text{RNA}}^{\text{mol}} N_{\text{RNA}} = 1.05 \times 10^{-2}$. RNA is highly negatively charged due to its phosphate backbone. Thus, we assign to each RNA bead a charge $q_{\text{RNA}} = -10\mathcal{Q}$, so that RNA molecules are uniformly charged with total charge $Q_{\text{RNA}} = -1000\mathcal{Q}$. The Ki-67 polymers are also uniformly charged (each bead having charge $q_0$), with the exception of the furthermost bead from the chromosome surface, which is given a higher charge $q_{\text{CP}}$ (Fig. 1A of the main text). The fully dephosphorylated state of Ki-67 (as found in mitotic exit) is modeled by choosing $q_{\text{CP}} = 24.15\mathcal{Q}$ and $q_0 = 3.72\mathcal{Q}$. The ratio $q_{\text{CP}}/q_0$ is chosen to approximate the corresponding experimental values, which were determined from the amino-acid sequence for the dephosphorylated state by assuming a total of 150 phosphorylation sites for the phosphorylated state [2]. To assess the role of the CP in the formation of RNA bridges between the two grafted surfaces, we additionally perform simulations in which the charge of the CP is uniformly spread over the rest of the Ki-67 molecules, so that the latter is uniformly charged and each bead has charge $q_{\text{unif}} = 4.57\mathcal{Q}$ (Fig. 1B of the main text). In both the CP and the uniform charge cases, the total charge of a Ki-67 molecule is $Q_{\text{Ki-67}} = 109.7\mathcal{Q}$.

The solvent is simulated implicitly using a Langevin thermostat, which also ensures that the temperature $T = 1$ is kept constant during the simulation. The viscous friction that each bead experiences is $\zeta = 10m/\tau$, so that the time it takes a free monomer to diffuse a distance $\sigma$ is $\tau_D = \sigma^2\zeta/k_BT = 10\tau$. The simulations are carried out using LAMMPS [8], and time integration is performed using the velocity Verlet algorithm, with time step $\delta t = 5 \times 10^{-3}\tau = 5 \times 10^{-4}\tau_D$. Periodic boundary conditions are applied in the $x$ and $y$ directions, *i.e.*, parallel to the grafted surfaces. Initially, the RNA polymers are arranged at random in the simulation box. After the overlaps are removed using a soft potential, the system is equilibrated for a time $\tau_{\text{equil}} = 2.5 \times 10^5\tau$ ($= 5 \times 10^7$ time steps) using only the excluded volume interactions Eq. (S1) before the production run with the full interactions. During production, which lasts for a time $\tau_{\text{prod}} = 10^6\tau$ ($= 2 \times 10^8$ time steps), we measure the $zz$-component of the pressure tensor, $P_{zz}$. This quantity corresponds to the force per unit area acting on each one of the two surfaces. We note that the pressure tensor is related to the stress tensor $\Sigma_{ij}$ by $P_{ij} = -\Sigma_{ij}/V$, where $V = L_zA$ is the system's volume [9], and that the $zz$-component of the pressure tensor is related to the total pressure $P$ of the system by $P = (P_{xx} + P_{yy} + P_{zz})/3$. Following the convention, the sign of this force is directed away from the middle of the box. Thus, if its value is negative, there is an attractive force acting between the two surfaces. If it is positive, the force is repulsive. For each set of parameters, we simulate five replicas of the system for the sparse brush and three replicas for the dense one, and average the results over the replicas.

#### C. Mapping to physical units

To draw a correspondence between simulation results and physical quantities, we map the simulation units of length and energy to SI units. The unit of energy in simulations is $\epsilon = k_BT$, where $k_B$ is Boltzmann's constant and $T$ the absolute temperature. Assuming $T = 310$ K (37°C), we have $k_BT \approx 4.3 \times 10^{-21}$ J ($\approx 2.6$ kJ/mol). For the length scale mapping, we can consider the height $\langle z \rangle_0$ of the unperturbed Ki-67 brush. In experiment, the measured brush height is $\approx 90$ nm in early mitosis (when Ki-67 is phosphorylated). In simulations, we can mimic phosphorylated Ki-67 by setting $q_{\text{CP}} = 6.17\mathcal{Q}$ (CP charge) and $q_0 = -8.54\mathcal{Q}$ (rest of the molecule charge) [2]. Under these conditions, the height of the unperturbed sparse brush with CP (grafting density $\rho_{\text{graft}} = 2.04 \times 10^{-2}\sigma^{-2}$) is $8.3\sigma$ (see Tab. S2). We thus impose $8.3\sigma = 90$ nm, which yields $\sigma = 11$ nm. The choice to use as reference the brush height in early mitosis is consistent with Hernandez-Armendariz *et al.* [2]. Moreover, we note that in the simulations the brush height only changes by $\approx 20\%$ between the dephosphorylated and phosphorylated state (see Sec. S3 A). Given this mapping, the simulation unit of charge is $\mathcal{Q} = \sqrt{4\pi\varepsilon_0\sigma\epsilon} = 7.3 \times 10^{-20}\text{C} = 0.45e$, with $e$ the elementary charge. The units of pressure and force are respectively $[P] = \epsilon/\sigma^3 = 3.2$ kPa and $[F] = \epsilon/\sigma = 0.40$ pN. The mapping between simulation and physical units is summarized in Tab. S1. We note, however, that this mapping should not be taken as exact, but rather as approximate, due to the coarse-grained nature of our minimal model.

### S2. ANALYTICAL MODEL

In this section, we present in detail the analytical model presented in the main text, and then discuss how it helps us shed light on the bridging mechanism observed in the simulations, and in particular on the role

| Physical quantity | Simulation unit | Experimental unit |
|---|---|---|
| Length | $\sigma$ | 11 nm |
| Energy | $\epsilon = k_B T$ | 2.6 kJ/mol |
| Charge | $\mathcal{Q} = \sqrt{4\pi\varepsilon_0\sigma\epsilon}$ | $7.3 \times 10^{-20}$ C |
| Pressure | $\epsilon/\sigma^3$ | 3.2 kPa |
| Force | $\epsilon/\sigma$ | 0.40 pN |

TABLE S1. Mapping between simulation units and physical units.

of the CP.

#### A. Flory-Huggins approach to coacervation

Here, we wish to illustrate why a uniformly charged brush causes RNA to fully mix with it, while a brush with a CP does not. To do so, we use the Flory-Huggins theory of polymer mixing [10]. We consider two plates separated by $L_z$ grafted with a uniformly charged polymer brush, in contact with a solvent containing RNA molecules occupying a volume fraction $\phi_2$. We use a lattice description where space is divided in sites of unit size occupied by RNA monomers, brush monomers, or solvent molecules. We assume that the brush occupies a layer of size $\langle z\rangle$ and we denote as $\phi_b$ the volume fraction of the brush there. The overall volume fraction of RNA (averaged over the brush and the solvent) is assumed to be constant and equal to $\phi_0$. For simplicity, we assume that $\langle z\rangle$ does not depend on $\phi_0$, and wish to compute the volume fraction $\phi_1$ of RNA within the brush at thermal equilibrium. The free energy of RNA per unit volume in the grafted region reads, in units of $k_{\mathrm{B}}T$:

$$f_1 = \frac{\phi_1}{N_{\mathrm{RNA}}}\log\phi_1 + (1-\phi_1-\phi_b)\log(1-\phi_1-\phi_b) - c\chi\phi_1\phi_b, \tag{S4}$$

where $N_{\mathrm{RNA}}$ is the number of beads in an RNA polymer, $\chi$ represents the enthalpic gain of a monomer in contact with the brush and $c$ is the lattice coordination number. Note that we treat the electrostatic interaction between the RNA and the brush as a contact interaction, as it is screened in the simulations. For the choice of interaction parameters chosen in the simulations of uniformly charged brushes, we have $\chi \approx 1$ and $c = 6$. The value $\chi = 1$ is obtained by evaluating Eq. (S2) for a RNA bead and a brush bead in contact. The free energy of RNA monomers in the solvent reads:

$$f_2 = \frac{\phi_2}{N_{\mathrm{RNA}}}\log\phi_2 + (1-\phi_2)\log(1-\phi_2). \tag{S5}$$

Here we do not consider any possible interaction between the two plates (*i.e.*, no RNA bridge). One can then minimize the total free energy $2\langle z\rangle f_1 + (L_z - 2\langle z\rangle)f_0$ numerically, while keeping the total number of RNA monomers constant so that $2\langle z\rangle\phi_1 + (L_z - 2\langle z\rangle)\phi_0 = N_{\mathrm{RNA}}n_{\mathrm{RNA}}$, with $n_{\mathrm{RNA}}$ the number of RNA polymers. We obtain that for $L_z \lesssim 400$, equilibrium is only possible when RNA is entirely depleted from the solvent and fully adsorbed onto the brush, as seen in simulations. RNA molecules start permeating the solvent for $L_z \approx 400$, when the brush saturates with RNA; the two plates are then too far to interact. This validates *a posteriori* the fact that bridging is negligible: for low $L_z$, RNA molecules are all adsorbed onto the brush.

Let us now carry out a similar derivation for the CP case. There are two main differences with the uniform case, due to the localized nature of CPs: (1) Since they are in limited number, one must account for potential saturation effects and (2) They are found only at the extremity of brush molecules. We crudely model this effect by assuming that the interaction between the brush and RNA only occurs in a thin layer, but corresponds to a higher energy per contact, $\chi' \approx 6$ (corresponding to a RNA bead in contact with a CP according to Eq. (S2)). We have now:

$$f_1 = \frac{\phi_1}{N_{\mathrm{RNA}}}\log\phi_1 + (1-\phi_1-\phi_b)\log(1-\phi_1-\phi_b) - c\chi'\phi_1\phi_{\mathrm{CP}}\left[1-\left(\frac{\phi_1}{c\phi_{\mathrm{CP}}}\right)^2\right], \tag{S6}$$

where $\phi_{\mathrm{CP}}$ is the volume fraction of CPs only. Here, the term in square brackets is a phenomenological term that prevents $\phi_1$ from reaching unphysically high values, and follows from the observation that the maximum value of $\phi_1$ that is physically possible is $c\phi_{\rm CP}$ (every single CP is bound to exactly $c$ RNA beads). After minimizing the total free energy while keeping the total RNA fraction $\phi_0$ constant, one finds that all RNA molecules stick to the brush only for $L_z \lesssim 20$; this values is however too low to safely neglect bridging (as it is of a similar order of magnitude as RNA's radius of gyration). Therefore, this simple Flory-Huggins approach fails to account for the phase behavior of RNA in presence of CPs.

In the next section, we will overcome this issue using a polymer model based on random walks and we will compute the bridging probability of RNA in both the uniform and the CP cases. We will find that bridging is essentially absent in the uniform case unless the two plates are in contact, and cannot be neglected in the CP case.

In what follows, an important parameter is the adsorption free energy of RNA beads into the brush. We assume that this free energy $\Delta F$ is equal to the excess free energy of adsorbed RNA beads in our Flory-Huggings theory. To obtain it, we consider a fictitious case where a single grafted plate would be at equilibrium with a solvent containing a fraction $\phi_0$ of RNA. The quantity $\phi_1$ of RNA adsorbed onto the brush can be obtained by equating the two chemical potentials:

$$\mu_1 = \frac{\partial f_1}{\partial \phi_1}(\phi_1) = \mu_0 = \frac{\partial f_2}{\partial \phi_2}(\phi_0), \tag{S7}$$

from which we obtain the excess free energy of adsorption as:

$$\Delta F = \log \frac{\phi_1}{\phi_0}. \tag{S8}$$

For $\phi_0 = 10^{-2}$ (corresponding to the volume fraction of RNA in simulations), we find the following values: CP dense, $\Delta F = 3.1$; CP sparse, $\Delta F = 2.3$; uniform dense, $\Delta F = 4.3$; uniform sparse, $\Delta F = 4.2$.

#### B. Model definition

The model is schematically represented in Fig. S2A. We assume that the two parallel plates are separated by a distance $L_z$, and bear a density $\rho_{\rm graft}$ of grafted polymer brush. The space between the plates is filled with RNA polymers. We denote by $\sigma = 1$ the size of polymer beads (monomers) and $\ell = 100$ the contour length of the RNA polymers. We additionally take $k_{\rm B}T$ as the unit of energy.

In the simulations, RNA is excluded from the vicinity of the plates in the CP case, due to steric repulsion with the polymer brush. To account for this effect in our model, we assume that RNA polymers can only explore the space from $x = 0$ to $x = D = L_z - 2h$, with $h$ the size of the excluded zone near each plate (located at $x = -h$ and $x = D + h = L_z - h$), as shown in Fig. S2B. In addition, we assume that RNA beads can adsorb into the brush with an free energy gain per bead $-\Delta F < 0$. This binding can occur in a region of thickness $\delta$ near each brush, *i.e.* $x \in [0, \delta]$ and $x \in [D - \delta, D]$. The values of $h$ and $\delta$ depend on whether the the brush is uniformly charged or has CPs, as described below.

For a dense brush with CPs, RNA is largely excluded from the brush so we assume that $h \approx \langle z\rangle$ the thickness of the brush. Binding only occurs when RNA beads are in close physical contact with CPs, so $\delta \approx 1$ (size of a bead), and we recall that, from our Flory-Huggins calculation, $\Delta F = 3.1$ (see Sec. S2 A). In the case of sparse brush with CP, the situation is similar: the binding strength with the CP is unchanged; however, RNA is not completely excluded from the brush. Instead, brush polymers adopt a mushroom-like configuration, where RNA is excluded from the mushrooms but can bind to the CPs anywhere on the surface of the mushrooms. We account for this intermediate phenomenology by setting $h = 4$ and $\delta = 3$ (so that $h + \delta = \langle z\rangle$ the average thickness of the brush). In this case $\Delta F = 2.3$.

In the uniform case, RNA can almost freely interpenetrate with the brush due to favorable electrostatic interactions. We will therefore assume that $h \approx 0$, and $\delta$ is comparable to the mean brush height $\langle z\rangle$. From the simulation data (Sec. S3 A), we therefore have $\delta \approx 7$ in the sparse case and $\delta \approx 9$ in the dense case. For the uniform brush, we have $\Delta F = 4.3$ in the dense case and $\Delta F = 4.2$ in the sparse case (Sec. S2 A).

#### C. Bridging probability

Below we derive an analytical expression for the bridging probability $p_{\rm bridge}$. We note that this is a different quantity from the fraction of bridging RNAs, as detailed in Sec. S2 D. The bridging probability is defined as

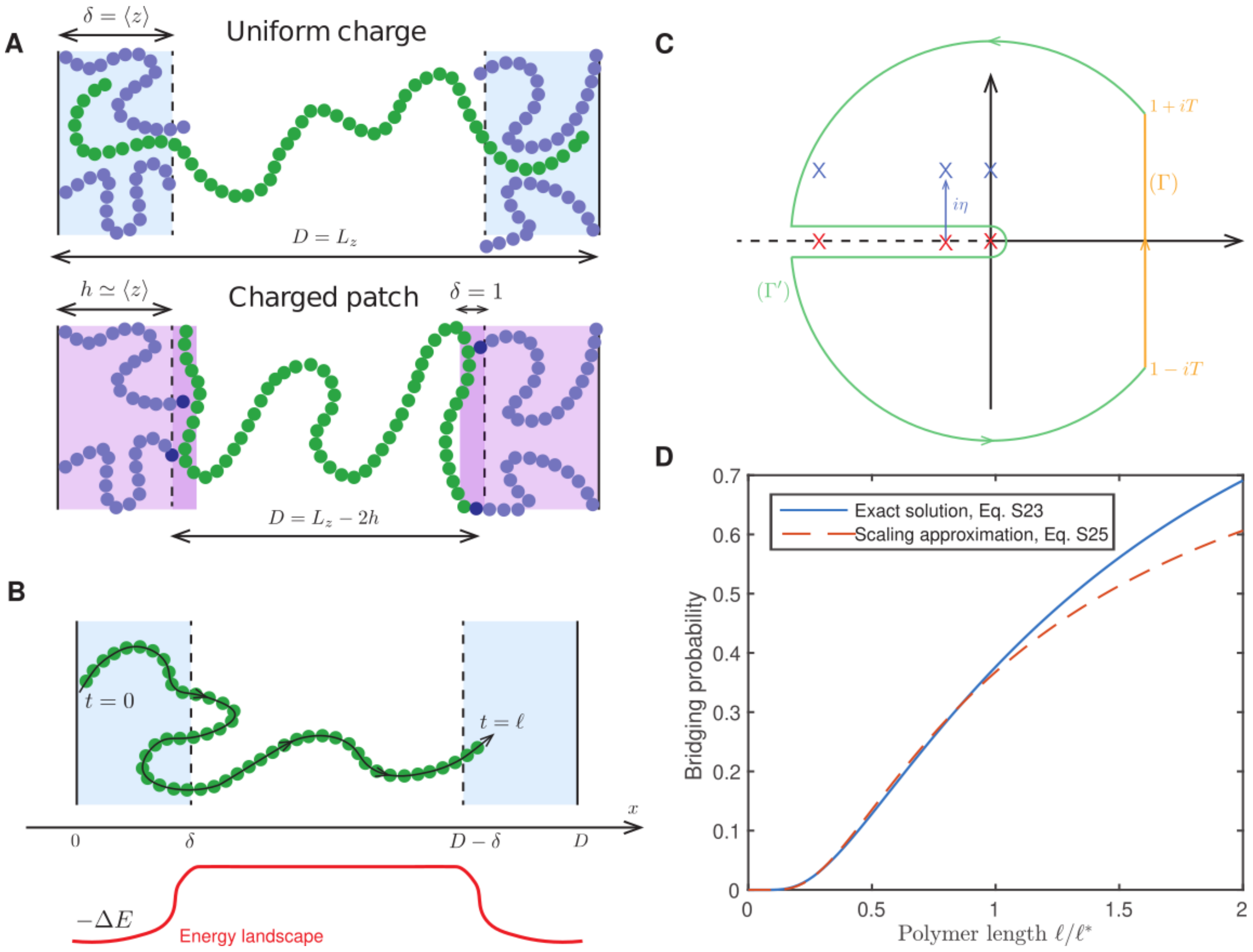


FIG. S2. Theoretical model of polymer bridging. **(A)** Schematic representation of the model. Uniformly charged brush (top): the bridging polymers can fully interpenetrate the grafted layer, so that $h = \langle z \rangle$. Brush with CPs (bottom): the bridging polymers do not penetrate the brush, so that $\delta = 1$ corresponds to the bead size. The polymers are therefore confined to a space $D = L_z - 2h \leq L_z$. **(B)** Generalized representation of the model. Bridging polymers (RNA) are modeled as ideal chains that can bind to a background of grafted polymers occupying a region of thickness $\delta$, in which binding can occur. For the uniform brush, $\delta \approx \langle z \rangle$ and $D \approx L_z$, whereas for the brush with CP, $\delta \approx 1$ and $D = L_z - 2h \approx L_z - 2\langle z \rangle$ (cfr. panel A). The variable $t$ represents the position along the polymer chain. **(C)** Integration contour for the inversion of the Laplace transform. The infinitely-many poles of the function $f$ are shifted into the upper complex half-plane by introducing a small shift $i\eta \to 0$. We close the integration path $(\Gamma)$ (in orange) by a key-hole shaped contour $(\Gamma')$ (in green). In the limit $T \to \infty$, the integral along $(\Gamma')$ vanishes. The integral along $(\Gamma)$ is then given by the residue theorem. **(D)** Probability $p_{\mathrm{bridge}}$ that a given polymer forms a bridge, as function of the rescaled polymer contour length $\ell/\ell^*$. Solid blue line: exact solution Eq. (S23). Dashed orange line: scaling approximation Eq. (S25).

the probability that an RNA polymer that is bound by one of its extremity to the surface located at $x = \delta$, reaches the other side. To compute this quantity, we use a continuous polymer model and define $p(\mathbf{x}, t)$ as the probability that the $t^{\mathrm{th}}$ monomer is located at position $\mathbf{x}$. Since we considered that the polymer behaves like an ideal chain, we can project the problem on the $x$ axis and work in 1D. The probability density $p$ is then the solution of a diffusion equation:

$$\partial_t p = \partial_{xx} p, \tag{S9}$$

where $t$ represents the position along the length of the polymer. The initial condition is $p(x, t=0) = \delta(x-\delta)$, with $\delta$ the Dirac-$\delta$ function, and one has the following boundary conditions:

$$p(x=\delta^-, t) = e^{\Delta F} p(x=\delta^+, t), \tag{S10}$$

$$\partial_x p(x=\delta^-, t) = \partial_x p(x=\delta^+, t), \tag{S11}$$

$$\partial_x p(x=0, t) = 0 \tag{S12}$$

$$p(x=D/2, t) = 0. \tag{S13}$$

where $x = \delta^+$ and $x = \delta^-$ are the limits $x \to \delta, x > \delta$ and $x \to \delta, x < \delta$, respectively. The equation $\partial_x p(x=0,t) = 0$ imposes a reflecting boundary condition at $x = 0$, whereas the last condition corresponds to an absorbing boundary condition at $x = D/2$, to match the definition of the bridging probability in numerical simulations. This is the probability that a polymer reaches the center of the simulation box – as polymers which do not are adsorbed onto the brush. In practice, we thus define the bridging probability $p_{\text{bridge}}$ as the probability that a particle that starts at $x = \delta$ at time $t = 0$ reaches the absorbing boundary condition at $x = D/2$ at some time before $t = \ell$ ($\ell$ being the contour length of the chain). This probability is equal to the time integral of the incoming probability flux at the absorbing boundary condition:

$$p_{\text{bridge}}(\ell) = \int_0^\ell -\partial_x p(D/2, t)\, \mathrm{d}t. \tag{S14}$$

We now introduce the Laplace transform defined as

$$\tilde{f}(s) = \int_0^{+\infty} e^{-st} f(t)\, \mathrm{d}t. \tag{S15}$$

In what follows we drop the tilde to lighten the notation.

The above system of equations can be solved in a straightforward manner in Laplace space, yielding:

$$p_{\text{bridge}}(s) = \frac{1}{s} \frac{2\cosh \delta\sqrt{s}}{(1+e^{\Delta F})\cosh \frac{D\sqrt{s}}{2} - (e^{\Delta F}-1))\cosh \frac{(D-4\delta)\sqrt{s}}{2}} = \frac{1}{sf(s)}, \tag{S16}$$

In what follows, we analyze this result in the two relevant limits ($D \gg \delta$, corresponding to the CP case, and $\delta \lesssim D$, corresponding to the uniform case). It should be noted that in both cases $D$ must be greater than $\delta$ for the model to hold (in other words, the polymer brush should not be too compressed). For values of $L_z$ such that $D$ would be smaller than $\delta$, we assume that $\delta = D/2$ (RNA can interact with the brush everywhere), which directly yields $n_{\text{bridge}} = 1$.

#### D. Charged patch case

In this case, $\delta \ll D$ and one can invert the above Laplace transform exactly as follows. Since $e^{\Delta F} \gg 1$, we find that

$$f(s) \approx \cosh D\sqrt{s}/2 + \alpha\sqrt{s}\sinh D\sqrt{s}/2. \tag{S17}$$

with $\alpha = \delta e^{\Delta F}$ (typically $\alpha \gg 1$). The function $n_{\text{bridge}}(s)$ possesses an infinite number of poles, all located on the negative real axis. We denote them as $(s_n)_n$, with $s_{-1} = 0$ and the $(s_n)_{n\geq 0}$ corresponding to the zeros of the function $f$ These zeros are obtained by solving the implicit equation

$$u_n = \cot \epsilon u_n, \tag{S18}$$

where $s_n = -u_n^2/\alpha^2$, $u_n \in [n\pi/\epsilon, (n+1)\pi/\epsilon]$ and $\epsilon = D/2\alpha$. For $n \geq 1$, $u_n$ approaches either $n\pi/\epsilon$ or $\pi(n+1/2)/\epsilon$ depending on the value of $\epsilon/n\pi$. For $n = 0$, $u_0$ approaches either $1/\sqrt{\epsilon}$ or $\pi/2\epsilon$ depending on the value of $\epsilon$. Overall, the following expressions provide a good approximation of the poles

$$u_0 \approx \frac{1}{\sqrt{\epsilon}} \frac{1}{\sqrt{1+4\epsilon/\pi^2}}, \tag{S19}$$

$$u_n \approx \frac{1}{\epsilon}\left[n\pi + \frac{\pi}{2} - \frac{\pi/2}{1+2\epsilon/n\pi^2}\right], \quad n \in \mathbb{N}^*. \tag{S20}$$

In particular, these expressions match the exact expansions of the $u_n$'s up to order 2 in $\epsilon$. In practice, we solve these equations numerically up to $n = 20$.

In addition to these poles, the function $f(s)$ also has a branch cut on $\mathbb{R}^-$. The inverse Laplace transform reads:

$$p_{\text{bridge}}(\ell) = \lim_{T\to\infty} \frac{1}{2i\pi} \int_{1-iT}^{1+iT} \frac{e^{s\ell}}{sf(s)}\,\mathrm{d}s, \tag{S21}$$

where the integration is performed on a vertical axis in the complex plane. Since poles are located on the branch cut, the integral cannot be evaluate through standards techniques involving the residue theorem. Instead, we use the following trick:

$$p_{\text{bridge}}(\ell) = \lim_{\eta\to 0} \frac{1}{2i\pi} \lim_{T\to\infty} \int_{1-iT}^{1+iT} \frac{e^{s\ell}}{sf(s) - i\eta}\,\mathrm{d}s, \tag{S22}$$

where the small parameter $\eta$ is used to evaluate the Cauchy principal value of the integral by shifting all poles in the upper half complex plane. We can then use a keyhole-shaped integration contour (see Fig. S2C) to evaluate the integral with the residue theorem. Integrals over the chosen contour are found to vanish in the limit $T \to \infty$, yielding:

$$p_{\text{bridge}}(\ell) \approx 1 + \sum_{n=0}^{\infty} \frac{e^{s_n\ell}}{s_n f'(s_n)}. \tag{S23}$$

This prediction is represented on Fig. S2D (blue solid line).

Alternatively, we can use a trick to estimate $n_{\text{bridge}}$ from the Laplace transform of its derivative:

$$p_{\text{bridge}}(\ell) = n_{\text{bridge}}(0) + \int_0^\ell \dot{p}_{\text{bridge}}(\tau)\,\mathrm{d}\tau \approx \int_0^\infty e^{-\tau/\ell}\dot{p}_{\text{bridge}}(\tau)\,\mathrm{d}\tau = \dot{p}_{\text{bridge}}(s = 1/\ell) = \frac{1}{f(1/\ell)}. \tag{S24}$$

This approximation suggests to use the following scaling expression:

$$p_{\text{bridge}}(\ell) \approx e^{-D^2/4\ell u^{*2}(\epsilon)} = e^{-\ell^*/\ell}, \tag{S25}$$

where the quantities $\ell^*$ and $u^*$ are defined by:

$$f(1/\ell^*) = \cosh u^* + \frac{1}{\epsilon} u^* \sinh u^* = 2. \tag{S26}$$

This prediction is represented on Fig. S2D (orange dashed line); it compares quantitatively to the exact solution. In particular, it reproduces the step-like shape of $n_{\text{bridge}}$ with the polymer length $\ell$. It shifts from $n_{\text{bridge}} = 0$ (no bridges) to $n_{\text{bridge}} = 1$ (all polymers form bridges) for a certain polymer length $\ell^* = D^2/u^*(\epsilon)$. In the limit $\epsilon = D/\alpha \to 0$, $u^* \approx \sqrt{D/\alpha}$, and in the limit $\epsilon \to \infty$ $u^*$ is a constant. For $D \gg \alpha \gg 1$, we have $\ell^* \approx D^2/2$, while for $\alpha \gg D \gg 1$ we have $\ell^* \approx D\alpha/2$.

Recall that to compare these results to numerical simulations, one should define $D$ as $L_z - 2h$ due to the excluded regions near the two plates. Lastly, we recall that $p_{\text{bridge}}$ is the probability that a given polymer that starts bound to a brush forms a bridge. The overall number of bridges is then given by:

$$n_{\text{bridge}} = n_{\text{RNA}} \times p_{\text{bound}} \times p_{\text{bridge}} \approx \frac{2\alpha n_{\text{RNA}}}{D + 2\alpha} e^{-\ell^*/\ell}, \tag{S27}$$

where $n_{\text{RNA}}$ is the number of RNA molecules and $p_{\text{bound}}$ the probability that a given polymer is bound to at least one brush. In Eq. (S27), we have assumed $p_{\text{bound}} \approx 1$, which is true for the simulation parameters used in the main text.

If the concentration of RNA is high enough to saturate the brushes (see also Sec. S3 C), we can as a first approximation replace the total number of RNA molecules $n_{\text{RNA}}$ in Eq. (S27) by the maximum number of RNA molecules that can bind to the brush – which we can approximate to the total number of CPs on each brush. A lower RNA concentration will not change the value of $p_{\text{bond}}$, as this is the probability that a given polymer is bound to at least one brush. Thus, for lower RNA concentrations we can expect Eq. (S27) to be valid, but with a lower $n_{\text{RNA}}$ value.

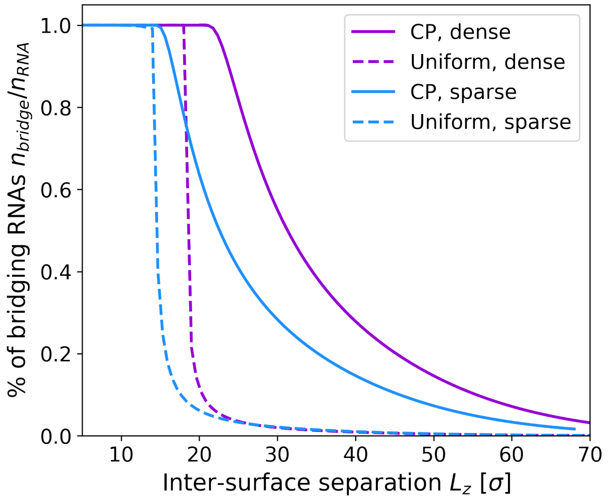


FIG. S3. Fraction of bridging RNAs $n_{\mathrm{bridge}}/n_{\mathrm{RNA}}$, Eq. (S27), for the different systems.

#### E. Scaling

The critical length $\ell^*$ can be expressed as function of the two parameters of the model, $\alpha$ (representing the strength of the binding between the brush and the polymers) and $D$ (the space available to polymers; for now we can assume that $D \approx L_z$ the physical distance between the plates):

$$\ell^* \approx \begin{cases} D^2/2 & \text{if} \quad D \gg \alpha \\ D\alpha/2 & \text{if} \quad D \ll \alpha \end{cases} \tag{S28}$$

The first case corresponds to a classical diffusion scaling.

Let us recover the $\ell^*$ scaling in the second case ($\alpha \gg D$) using scaling arguments. Each time the polymer touches one plate, an average of $\alpha$ monomers stay adsorbed on it. Each time the polymer leaves one plate, it has a probability $1/D$ of reaching the opposite plate without coming back to its starting position; this first-passage process therefore arrives once every $D$ "attempts". If the polymer reaches the other plate exactly once, then on average it will have left $\alpha D$ monomers on the first plate ($\alpha$ at each of the $D$ attempts). If this quantity is larger than the total number $\ell$ of monomers in the polymer, then bridging is impossible. Hence, bridging only occurs if $\ell > \ell^* = D\alpha$.

#### F. Uniformly charged brush

In the case of the uniformly charged brush, $\delta$ is large and one cannot make the approximation $\delta \ll D$. Instead, we numerically compute the inverse Laplace transform using the Talbot algorithm. In Fig. 4 of the main text, where $n_{\mathrm{bridge}}$ is plotted for all possible cases, we compute all inverse Laplace transforms using the Talbot algorithm.

#### G. Results and discussion

In Fig. S3 we plot the fraction of bridging RNAs $n_{\mathrm{bridge}}/n_{\mathrm{RNA}}$ for the different systems. The main take-away of the analytical model here presented is that, in the uniformly charged case, it is strongly favorable for RNA to be adsorbed onto the brush. This results is only weak bridging. On the other hand, in the CP case, the brush sterically excludes RNA, which favors bridging. In addition, while the binding of individual RNA beads is stronger with the CP than with beads from the uniformly charged brush, the number of beads that can actually bind to a single CP is limited due to steric and electrostatic repulsion between RNA beads, resulting in moderate adhesion compared to the uniform case.

Overall, the distance over which RNA bridges can form depends on the quantity $\alpha = \delta e^{\Delta F}$, where we recall that $\Delta F$ is the adsorption free energy and $\delta$ is the distance over which RNA penetrates the brush. In the case of the CP brush we have $\alpha \approx 10$, corresponding to a partial coacervation regime and a high bridging probability. On the other hand, the uniform brush corresponds to $\alpha \approx 100$, implying a complete coacervation and a low bridging probability.

### S3. ADDITIONAL SIMULATION DATA

In the following sections, we report additional simulation data, and additionally check the robustness of our results upon varying several parameters.

#### A. Unperturbed brush height

In experiment, the measured brush height is $\approx 90$ nm in early mitosis (when Ki-67 is phosphorylated) [2, 11]. The grafting density of Ki-67 on chromosomes is $\rho_{\text{graft}} \approx 210$ molecules μm$^{-2}$ [11]. In simulations, phosphorylated Ki-67 can be modeled by setting $q_{\text{CP}} = 6.17\mathcal{Q}$ (CP charge), $q_0 = -8.54\mathcal{Q}$ (rest of the molecule charge) for the CP and $q_{\text{unif}} = -7.93\mathcal{Q}$ for uniform charge [2]. In both cases, the total charge of a Ki-67 molecule is $Q_{\text{Ki67}} = -190.3\mathcal{Q}$. We measured the unperturbed height of the Ki-67 brush by simulating a single Ki-67 brush without RNA. We considered both sparse (grafting density $\rho_{\text{graft}} = 2.04 \times 10^{-2}\sigma^{-2}$) and dense ($\rho_{\text{graft}} = 6.25 \times 10^{-2}\sigma^{-2}$) brushes.

The results, reported in Tab. S2,can be summarized as follows: (i) The brush height is $\approx 30\%$ larger for the dense brush than for the sparse brush. This increase of brush height with grafting density is qualitatively expected for polymer brushes [6]. (ii) The brush height is very slightly larger for the uniformly charged brush than for the one with CPs. This is due to the on average higher electrostatic repulsion between the Ki-67 beads in the uniform case. (iii) The brush height is $\approx 20\%$ lower for the dephosphorylated brush than for the phosphorylated one. This is in qualitative agreement with experimental measurements [2, 11], which were however performed in the presence of RNA. The behavior of the brush height in the presence of RNA is discussed in Sec. S3 B below.

| **Phosphorylation state** | **Grafting density** | **CP/Uniform** | **Brush height $\langle z \rangle_0/\sigma$** |
|---|---|---|---|
| Dephosphorylated* | Sparse | CP | $7.2 \pm 0.2$ |
| Dephosphorylated* | Sparse | Uniform | $7.3 \pm 0.2$ |
| Phosphorylated | Sparse | CP | $8.3 \pm 0.2$ |
| Phosphorylated | Sparse | Uniform | $8.6 \pm 0.3$ |
| Dephosphorylated* | Dense | CP | $9.2 \pm 0.2$ |
| Dephosphorylated* | Dense | Uniform | $9.4 \pm 0.2$ |
| Phosphorylated | Dense | CP | $11.0 \pm 0.3$ |
| Phosphorylated | Dense | Uniform | $11.3 \pm 0.3$ |

TABLE S2. Brush height for brushes with different grafting densities (sparse and dense) and different Ki-67 charge distributions (CP and uniform). The * highlights the phosphorylation state used throughout this work.

#### B. Brush height and force between the two grafted surfaces

In Fig. S4A, we show the mean brush height $\langle z \rangle$, normalized by the height of the unperturbed brush $\langle z \rangle_0$, measured by simulating a single Ki-67 brush without RNA (see Sec. S3 A. The mean brush height is shown as a function of the normalized inter-surface separation $L_z/2\langle z \rangle_0$. For small values of the inter-surface separation, the two brushes interpenetrate ($L_z/2\langle z \rangle_0 \lesssim 1$). In this regime, the brush height increases approximately linearly as the distance between the surfaces increases. For $L_z/2\langle z \rangle_0 \approx 1 - 1.3$, the brush height reaches a maximum: this corresponds roughly to the regime of maximum bridging. Here, the brushes do not interpenetrate significantly, and, at the same time, the RNA bridges favor the extension of the Ki-67 molecules. As $L_z$ is further increased, we observe a monotonic decrease in the case of the dense brush, as the RNA is partially adsorbed onto the brush but mainly interacts with the CPs. For the sparse brush, on the other hand, the brush height displays an interesting non-monotonic behavior. We attribute this to the RNA being completely adsorbed onto the brush, resulting in partial saturation of the brush and a consequent extension of the brush polymers.

In Fig. S4B, we show the force $F_z$ acting between the two surfaces. The potential of mean force (PMF, see main text) is derived by integrating this quantity. As for the PMF, we normalize here the force by the

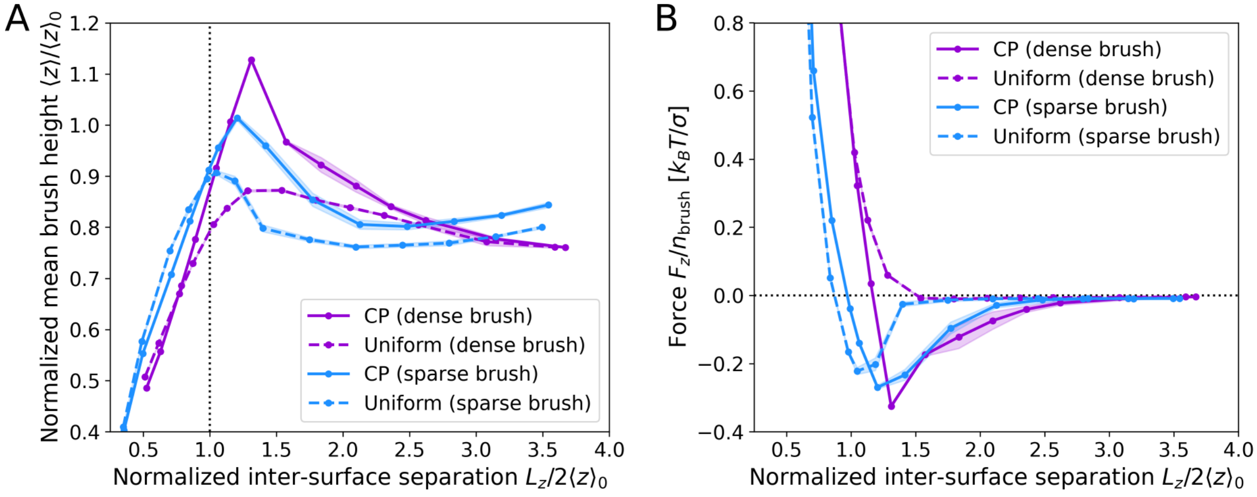


FIG. S4. **(A)** Mean brush height $\langle z \rangle$ as a function of the inter-surface separation $L_z$. Both the horizontal and the vertical axes were normalized by the mean height of the unperturbed brush $\langle z \rangle_0$. The values of $\langle z \rangle_0$ for the different systems are reported in Tab. S2. The maximum of $\langle z \rangle$ corresponds approximately to the minimum of the potential of mean force (Fig. 2 of the main text) and of the force acting between the two surfaces. **(B)** Force acting between the two grafted surfaces, computed as $F_z = AP_{zz}$, with $A$ the surface area and $P_{zz}$ the $zz$-component of the pressure tensor. The force is normalized by the number of polymers per brush $n_{\mathrm{brush}}$.

number of Ki-67 molecules per brush. By comparing Fig. S4B and Fig. S4A, we see that the minimum of $F_z$, corresponding to the largest attractive force, corresponds to the maximum brush extension. This confirms the intuition that the brush becomes maximally extended as a consequence of the formation of RNA bridges that pull the two surfaces together.

#### C. Effect of increasing RNA concentration

In this Section, we study the effect of increasing the RNA concentration, in particular to investigate whether saturation of the brushes by RNA could suppress the inter-surface attraction. In order to do so, we have performed simulations where the RNA density is double the one used in the main text. In these simulations, the density of RNA molecules is thus $\rho^{\mathrm{mol}}_{\mathrm{RNA}} = 4 \times 10^{-4}\sigma^{-3}$ instead of $\rho^{\mathrm{mol}}_{\mathrm{RNA}} = 2 \times 10^{-4}\sigma^{-3}$; the corresponding values of the RNA volume fractions are respectively 2% and 1%.

In Fig. S5A-B we show the force between the grafted surfaces $F_z$, normalized by the number of polymers per brush $n_{\mathrm{brush}}$, as a function of the normalized inter-surface separation, comparing the low and high RNA density cases. For the uniform dense brush (Fig. S5A, dashed lines), $F_z$ remains very similar when increasing RNA density, with the repulsive part being slightly shifted to lower inter-surface separation. This suggests that this RNA concentration is insufficient to saturate the uniform dense brush. In the CP case (Fig. S5A, solid lines), on the other hand, we observe a reduction of the attractive force ($F_z < 0$ region). We attribute this to the saturation of the CPs: due to the larger concentration of RNA molecules, single RNA molecules are more likely to bind to a single brush, saturating the CPs and leading to the formation of fewer bridges.

Interestingly, for the sparse brush, we observe a rather different behavior (Fig. S5B). Both for the uniform brush (dashed lines) and for the one with the CPs (solid lines), the attraction between the surfaces increases when increasing the RNA concentration. This effect is not due to the formation of a larger number of bridges, as we show in Fig. S6 and discuss below. Instead, the higher RNA concentration leads to a reduction of the repulsion between the positively charged brushes, which in turn leads to an effectively stronger attraction. This effect is also present for the dense brush, but in this case the repulsion between the brushes remains high due to the steric contribution resulting from excluded volume interactions.

We also observe an increase of $F_z$ at large inter-surface separation for high RNA concentration. This is an artifact of our simulation setup, where we only simulate one side of the chromosome surface, and thus the RNA molecules which are not adsorbed onto the brush exert a positive pressure between the surfaces. In reality, this pressure will be canceled out by an equal and opposite pressure exerted on the other side of the chromosome.

Due to this artifact, which only appears when there are free (*i.e.*, non-adsorbed and non-bridging) RNA

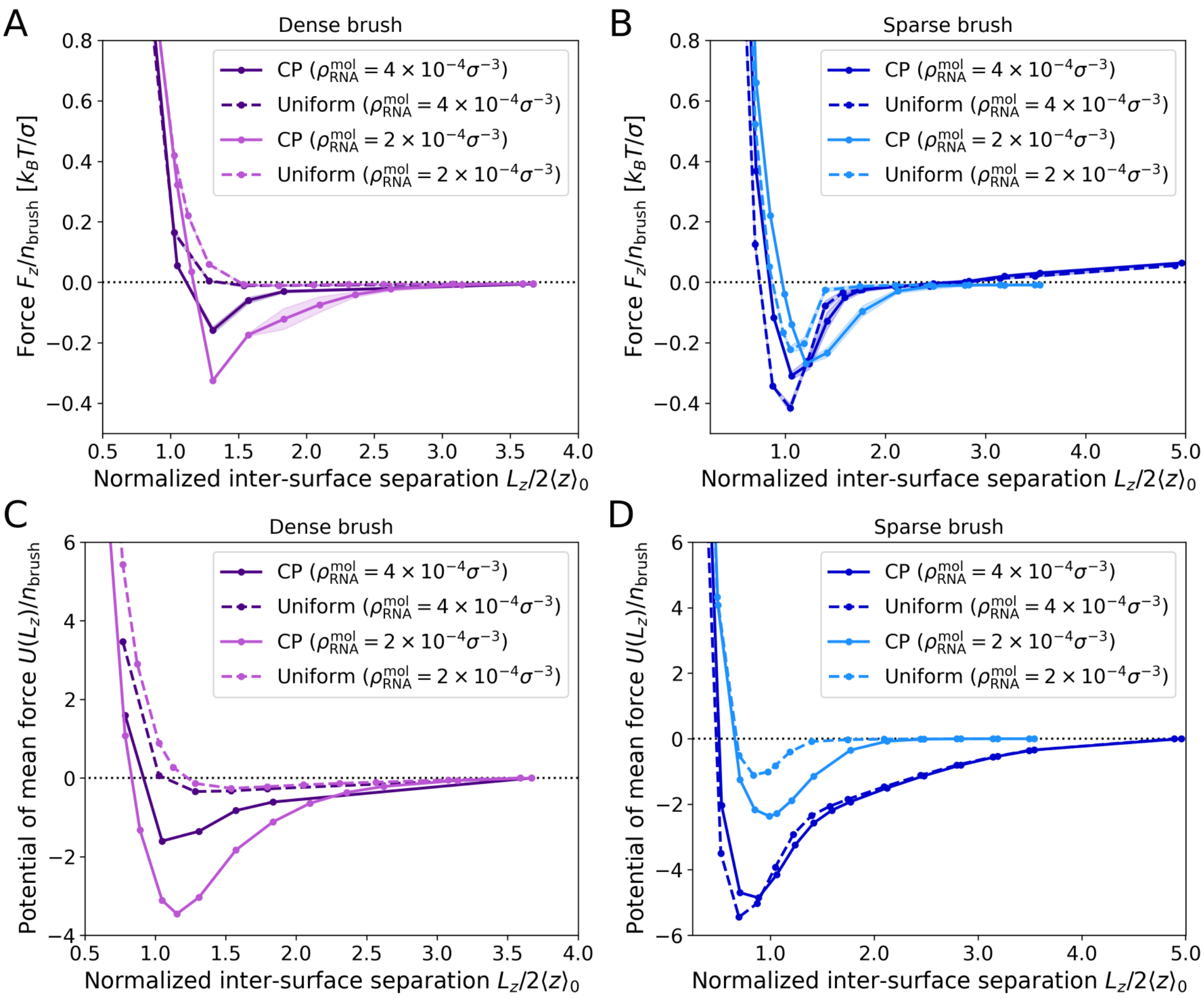


FIG. S5. **(A,B)**: Force (in the $z$ direction) acting between the two grafted surfaces, normalized by the number of polymers per brush $n_{\rm brush}$, for different RNA concentrations: $\rho_{\rm RNA}^{\rm mol} = 4 \times 10^{-4}\sigma^{-3}$ (darker colors) and $\rho_{\rm RNA}^{\rm mol} = 2 \times 10^{-4}\sigma^{-3}$ (lighter colors – value used in the main text). **(A)**: Dense brush. **(B)**: Sparse brush. **(C,D)**: PMF (see Eq. (S29)) between the surfaces. **(C)**: Dense brush. **(D)**: Sparse brush.

molecules for large $L_z$ values, one has to modify the definition of the PMF to remove the finite pressure at $L_z \to \infty$. Instead of using the definition of Eq. (1) of the main text, we define

$$U(L_z) \equiv \int_{L_z}^{\infty} [F_z(z) - F_z^{\infty}]\, {\rm d}z = A \int_{L_z}^{\infty} [P_{zz}(z) - P_{zz}^{\infty}]\, {\rm d}z, \tag{S29}$$

with $A$ the surface area, so that the infinite-distance contribution is removed. In Figs S5C-D, we compare the PMF at low and high RNA density using the definition (S29). We note for the lower RNA density used in the main text, this definition is equivalent to the original one, as $P_{zz}^{\infty} = 0$.

In Fig. S6, we show the fraction of bridging RNAs as a function of the normalized inter-surface separation for different RNA concentrations. One can see that for the dense brush with CP (A, solid lines), increasing the RNA concentration indeed leads to the formation of more bridges. For the sparse brush (B), instead, increasing the RNA concentration leads to the formation of the same amount of bridges in the uniform case (dashed lines), and to fewer bridges in the CP case (solid lines). We thus conclude that the stronger attraction observed for the sparse brush at higher RNA concentration (Fig. S5B and D) is indeed due to a reduction of the electrostatic repulsion between the brushes.

We note that to compute the number of bridges, we have used a slightly different definition from the one used in the main text (see *definition 2* in Sec. S3 F). This is because the original definition (a bridge is an

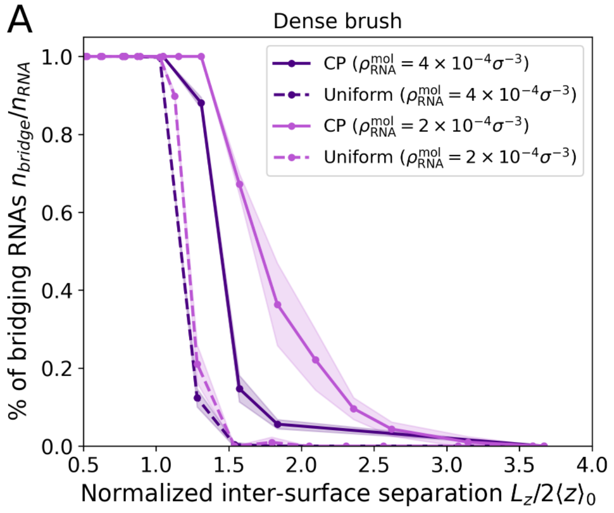


FIG. S6. Fraction of bridging RNAs as a function of the normalized inter-surface separation for different RNA concentrations: $\rho^{\rm mol}_{\rm RNA} = 4 \times 10^{-4}\sigma^{-3}$ (darker colors) and $\rho^{\rm mol}_{\rm RNA} = 2 \times 10^{-4}\sigma^{-3}$ (lighter colors – value used in the main text)

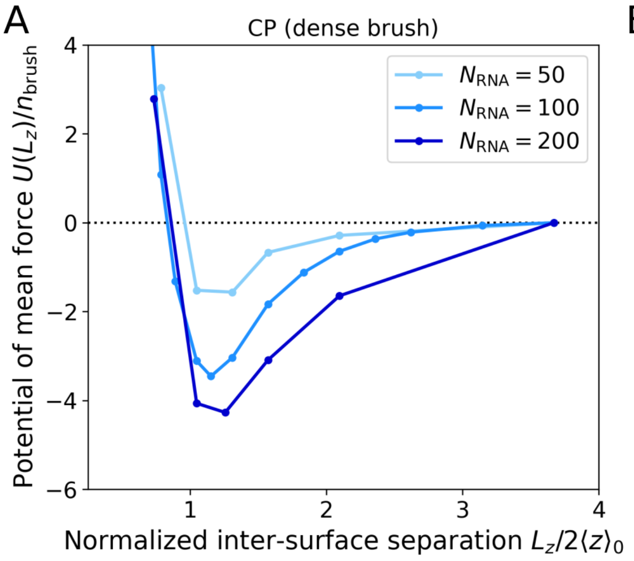


FIG. S7. Potential of mean force (PMF) between the grafted surfaces for the dense brush and for different RNA lengths $N_{\rm RNA}$, at constant RNA concentration matching the one used in the main text. **(A)**: CP case. **(B)**: Uniform case.

polymer whose end monomers are neither both in the portion $0 < z < L_z/2$ of the box nor in $L_z/2 < z < L_z$) does not work well in the presence of free RNAs at large $L_z$, since several RNAs that are free floating in the middle of the box will be classified as bridging.

To summarize, increasing the RNA concentration has two effects: In the CP case, it can lead to the saturation of the CPs and to the formation of a smaller number of bridges, resulting in a reduced attraction (Figs. S5A and C and S6A). Additionally, having more RNA adsorbed onto the brush reduces (screens) the electrostatic repulsion between the positively charged brushes, effectively causing an increase in the attraction. The total attraction strength thus results from these competing effects.

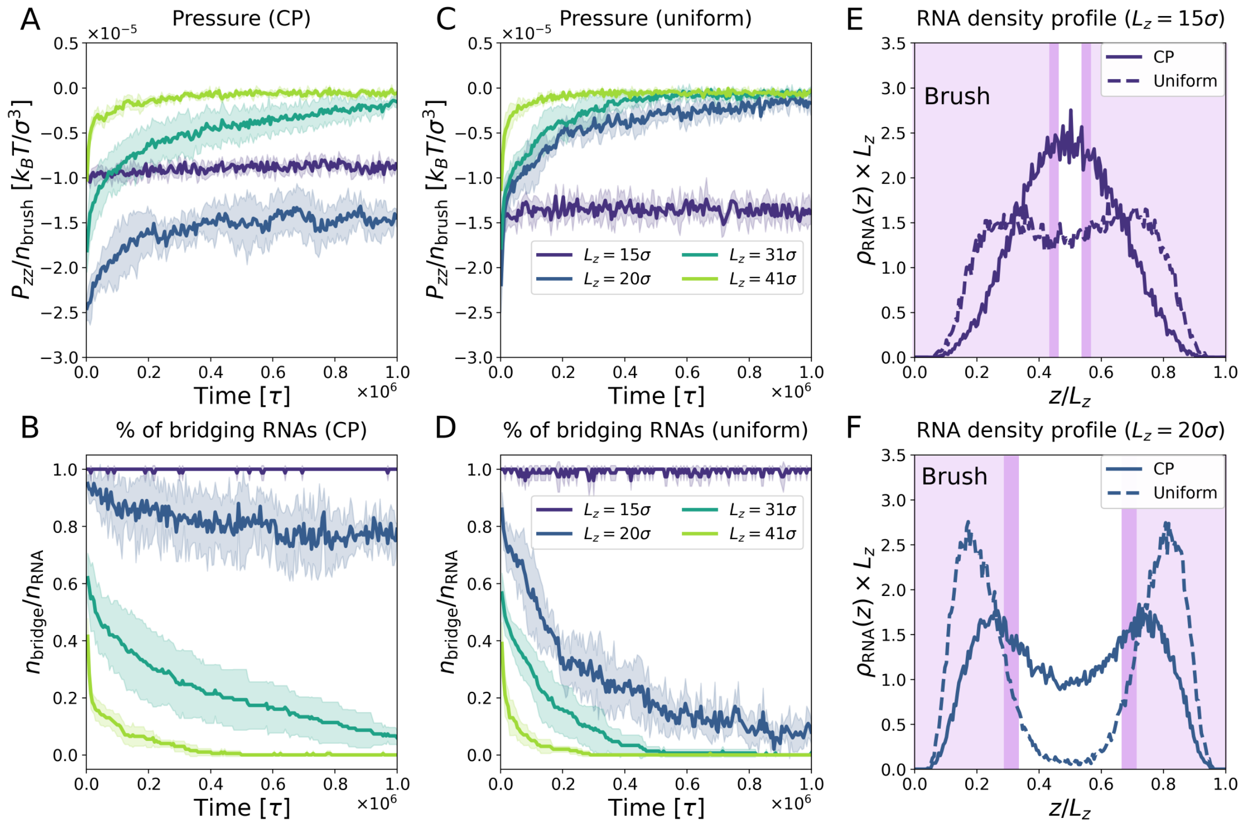


FIG. S8. **(A,C)** Time evolution of the $zz$-component of the pressure tensor $P_{zz}$, normalized by the number of grafted polymers per brush $n_{\mathrm{brush}}$. **(A)** CP, sparse. **(C)** Uniform, sparse. **(B,D)** Time evolution of the fraction of bridging RNAs, $n_{\mathrm{bridge}}/n_{\mathrm{RNA}}$. **(B)** CP, sparse. **(D)** Uniform, sparse. **(E-F)** Normalized $z$-density profile of RNA for the dense brush with inter-surface separation $L_z = 15\sigma$ **(E)** and $L_z = 20\sigma$ **(F)**. The shaded area represents the extension of the brush, with the lighter shade for the uniformly charged brush and the darker shade for the CP one.

#### D. Effect of changing RNA length

Throughout this work, we have modeled RNA as a 100-bead polymer. However, the size of premature ribosomal RNA varies. In this Section, we thus address the effect of changing the length of the RNA molecules. To this end, we performed simulations for the dense brush case for lengths $N_{\mathrm{RNA}} = 50$ and 200, corresponding respectively to half and double the length used in the main text. The total RNA density is kept constant and equal to the one used in the paper. As the Rouse relaxation time of RNA scales as $N_{\mathrm{RNA}}^2$, for $N_{\mathrm{RNA}} = 200$ we run the simulations for $4\times$ longer than for $N_{\mathrm{RNA}} = 100$ before measuring the PMF: this is needed in order to give enough time to the bridges to be released.

The PMF for the different lengths is reported in Fig. S7 for the CP (A) and uniform (B) cases. In the CP case (Fig. S7A), increasing $N_{\mathrm{RNA}}$ leads to an increase of both the range and the depth of the attractive well of the PMF. Thus, as expected, longer polymers lead to stronger and more long-range attraction. For the uniformly charged brush, no significant difference is observed between $N_{\mathrm{RNA}} = 50$ and 100. However, for $N_{\mathrm{RNA}} = 200$, the interaction becomes more attractive and long-ranged. We attribute this to the fact that a very long chain needs to be stretched less (compared to its length) to bridge between the two surfaces, *i.e.*, the relative entropic cost of forming a bridge is smaller. The strength and range of the interaction, however, remains significantly larger in the CP case also for $N_{\mathrm{RNA}} = 200$.

#### E. RNA bridging for the sparse brush

In Fig. S8, we report the time-evolution of the $zz$-component of the pressure tensor (A,C) and of the fraction of bridging RNAs (B,D) for the sparse brush, for selected values of the inter-surface separation $L_z$. In Fig. S8E,F, we report the density profile of RNA along the $z$ axis for the CP (solid lines) and uniform

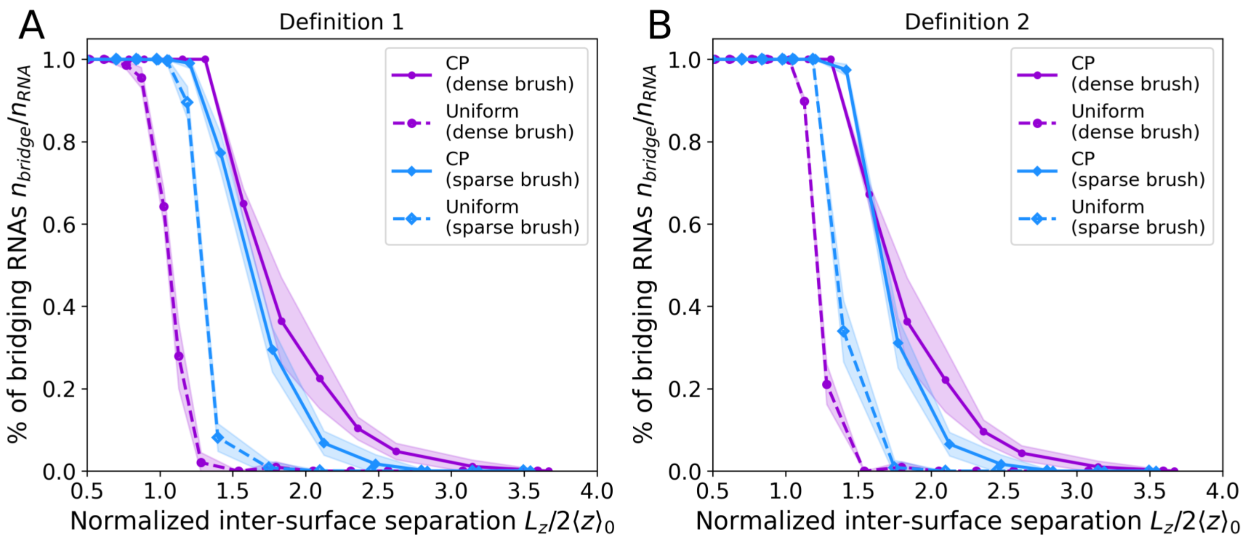


FIG. S9. Fraction of bridging RNAs as a function of the normalized inter-surface separation. Here, we compare two different definitions of bridging RNA (see text). **(A)**: definition 1 (used throughout this work). **(B)**: definition 2.

(dashed line) cases, for $L_z = 15\sigma$ (E) and $L_z = 20\sigma$ (F). As for the dense brush, we observe that the RNA molecules localize more towards the center of the box in the presence of the CP than with the uniformly charged brush.

#### F. Alternative definition of bridge

In this Section, we compare the definition of bridge used in the main text to an alternative one, showing that they give the same results in the range of parameters considered.

In the main text, we defined a bridge as (**definition 1**) any RNA polymer whose end monomers are neither both in the portion $0 < z < L_z/2$ of the box nor in $L_z/2 < z < L_z$. This definition is well suited to the case in which all RNA polymers are either bridging or adsorbed onto the brush, which is usually the case in our simulations. From visual inspection, it turned out to be the one that most reliably captures the correct number of bridges in our simulations.

However, in cases in which there is a significant fraction of free RNA polymers (*i.e.*, that are neither bridging nor adsorbed, see Sec. S3 C), or partially adsorbed RNA polymers (*i.e.*, that are attached to one brush but protrude significantly from it), another definition might be more suitable. We thus introduce the **definition 2**: a bridge is any RNA polymer that has one end bead in the volume between $z = 0$ and $z = z_{\max}^{\text{low}} + \sigma$ and the other end bead between $z = L_z - (z_{\max}^{\text{up}} + \sigma)$ and $z = L_z$, with $z_{\max}^{\text{up,low}}$ the maximal extension of polymers in the upper and lower brush in the given configuration, respectively.

In Fig. S9, we compare the two definitions for the systems investigated in the main text. As expected, the two definitions are equivalent at large inter-surface separations, although they differ slightly when the two brushes are close to each other ($L_z/2\langle z\rangle_0 \approx 1$).

#### G. Decay of number of bridges in time

In Fig. S10 we show the time evolution of the number of bridging RNAs on a semi-logarithmic scale for different systems. We recall that a bridge is defined as any RNA polymer whose end monomers are neither both in the portion $0 < z < L_z/2$ of the box nor in $L_z/2 < z < L_z$. For all systems but the dense brush with CP one (D), the number of bridges initially drops quickly as the electrostatic interactions are turned on and a fraction of RNA molecules becomes adsorbed onto the brushes. After this short-time transient, the number of bridges decreases approximately exponentially in time, *i.e.* $n_{\text{bridge}}(t)/n_{\text{bridge}}(0) \propto e^{-t/\tau_b}$, with $\tau_b$ a typical timescale for bond breaking. This exponential decay is what is expected if all bridges have approximately the same binding energy $E$ (so that the rate of bond breaking is $k \propto \tau_b^{-1} \propto e^{-E/k_BT}$) and if they break randomly and independently. The non-exponential decay observed for the dense brush with CP is likely due

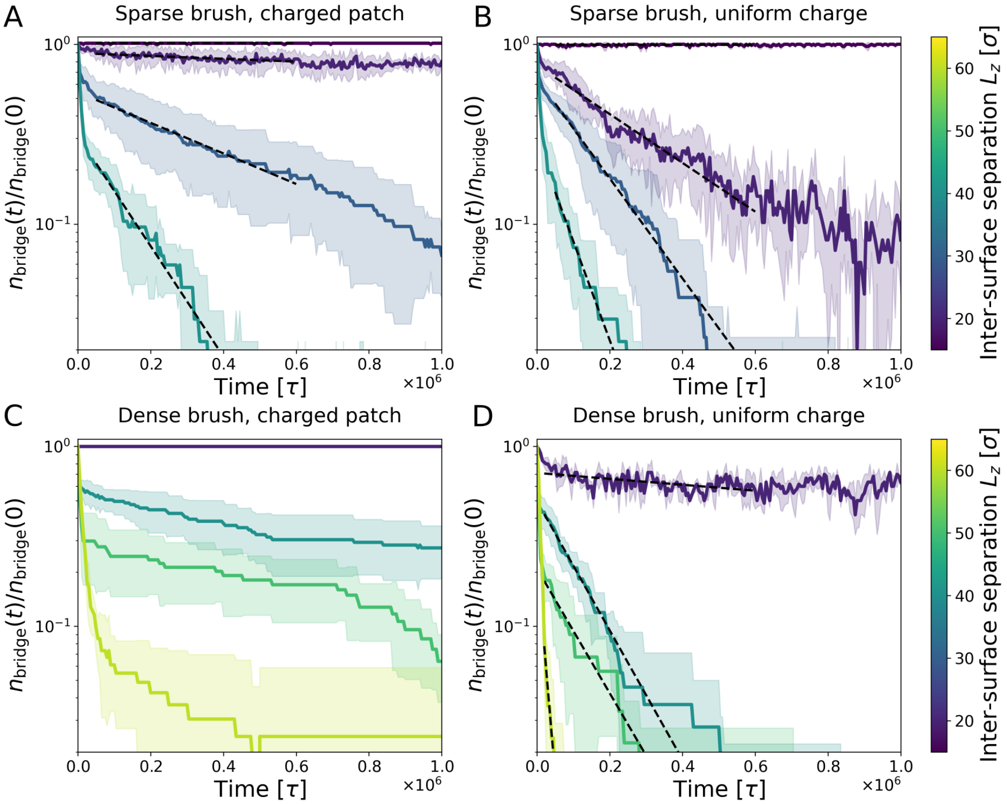


FIG. S10. Number of bridging RNAs as a function of time, normalized by the $t = 0$ value. **(A)**: Sparse brush, CP. **(B)** Sparse brush, uniform. **(C)** Dense brush, CP. **(D)** Dense brush, uniform. In **(A,B)** and **(D)**, the decay of the number of brushes is approximately exponential, as one might expect for a random bonding process. In **(B)**, the decay is non-exponential, signaling the possible presence of a wide distribution of binding energy for the bridges. Dashed lines are fits with the function $Ae^{-t/\tau_b}$, with $A < 1$ a positive constant. In A and B, the $t$-range for the fit is $5 \times 10^4\tau \leq t \leq 6 \times 10^5\tau$. In D, we considered a slightly different range, $2 \times 10^4\tau \leq t \leq 6 \times 10^5\tau$. This was done to exclude both the short-time transient and the noisy long-time regime.

to the fact that each RNA binds to multiple CPs, resulting in a wide distribution of binding energies. These different binding energy might lead to several exponential relaxation processes, which would superimpose to give a non-exponential relaxation [12].

#### H. Density profiles of RNA, brush monomers and CP along the $z$ axis

We report here the density profile along the $z$ axis of the RNA molecules (Fig. S11), of the brush molecules –excluding the CP– (Fig. S12), and of the CPs (Fig. S12). In all cases, the density profile was averaged over the time window $t \in [0.975, 1.00] \times 10^6\tau$, with $10^6\tau$ corresponding to the longest simulated time. Thus, we report the long-time average of the density profile. Also, in all plots the horizontal axis is normalized by the inter-surface separation $L_z$; the vertical axis is multiplied by $L_z$ so that the integral of the density profile is equal to 1. The data are shown for a range of inter-surface separations $L_z$ ranging from interpenetrating brushes (small $L_z$) to completely separated brushes (large $L_z$).

In Fig. S11, we show the density profile of the RNA molecules. By comparing the CP case (A: sparse brush; C: dense brush) to the case of uniformly charged brush (B: sparse brush; D: dense brush), one can clearly see that in the presence of the CP the RNA molecules localize in the middle of the box ($z/L_z$=0.5) much more strongly than with the uniformly charged brush, so that there is much higher bridging in this case. One can also see that in the uniformly charged brush case, there is a sudden transition to bridging when the two brushes are in contact with each other to no bridging as soon as they are separated. In the latter case, most of the RNA molecules are adsorbed onto the brush.

The density profile of the brush polymers is reported in Fig. S12. Overall, these density profiles correlate with those of the RNA, in that for the CP systems (A,C) there is a higher concentration of Ki-67 towards

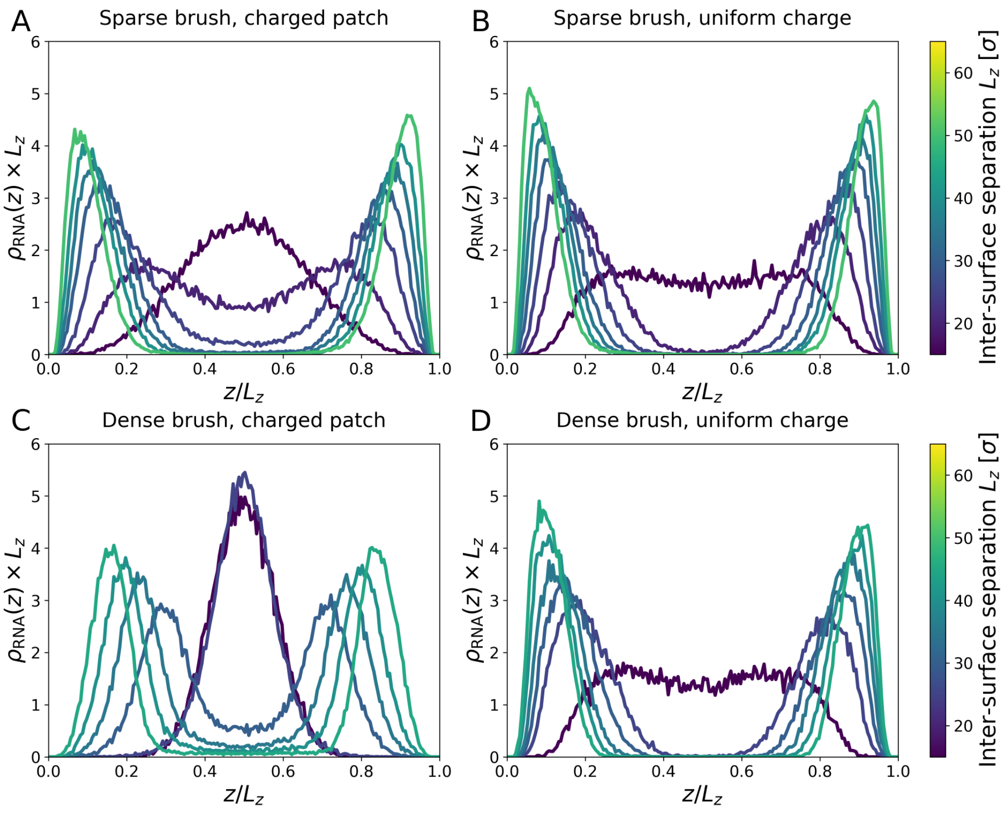


FIG. S11. Density profile of RNA along the $z$ axis. The horizontal axis was normalized by the inter-surface separation $L_z$. **(A)** Sparse brush, CP. **(B)** Sparse brush, uniform charge. **(C)** Dense brush, CP. **(D)** Dense brush, uniform.

the middle of the box, due to the RNA-mediated bridging. When the brush is uniformly charged (B,D), the brush collapses due to the coacervation of Ki-67 and RNA (compare with Fig. S4A).

Finally, in Fig. S13 we show the density distribution of the CPs for sparse **(A)** and dense **(B)** brushes. In the dense brush, the Ki-67 molecules adopt a more extended configuration due to the excluded volume interaction with their neighbors. As a result, the CPs are more strongly localized towards the center of the box.

### I. PMF dependence on brush polymer charge (dense uniform case)

When comparing the uniformly charged brush to the brush with the CP, we have imposed that the total charge of each brush polymer must be the same. Another approach consists in measuring the average electrostatic interaction energy $\langle E_{\mathrm{el,RNA}}\rangle$ of each bead of an RNA polymer interacting with the brush, and setting this quantity to a constant instead. In the case of dense brush with CP, we measured $\langle E_{\mathrm{el,RNA}}\rangle = (-4.97 \pm 0.02)k_BT$, whereas for the uniformly charged brush one has $\langle E_{\mathrm{el,RNA}}\rangle = (-2.99 \pm 0.02)k_BT$ (not considering the electrostatic interactions between RNA beads). The charge of each bead of a brush polymer in the uniform case is $q_{\mathrm{unif}} = 4.57\mathcal{Q}$. To compare the CP case to cases with similar values of $\langle E_{\mathrm{el,RNA}}\rangle$, we considered uniformly charged brushes with $q_{\mathrm{unif}} = 6.5\mathcal{Q}$, corresponding to $\langle E_{\mathrm{el,RNA}}\rangle = (-5.56 \pm 0.04)k_BT$, and with $q_{\mathrm{unif}} = 7.5\mathcal{Q}$, corresponding to $\langle E_{\mathrm{el,RNA}}\rangle = (-6.87 \pm 0.08)k_BT$. The corresponding total interaction potential, sum of the excluded volume and electrostatic components,

In Fig. S14B, we compare the potential of mean force (PMF, see Fig 2 in the main text) for these three cases of dense, uniformly charged brush with different values of the brush bead charge $q_{\mathrm{unif}}$. Upon increasing $q_{\mathrm{unif}}$, due to stronger electrostatic repulsion between the interpenetrating brushes, the repulsive part of the potential becomes steeper. However, the depth of the shallow attractive well remains roughly unchanged, and the range of the attractive region is reduced. This is due to the fact that for higher brush charge, the RNA polymer are adsorbed even more strongly into the brush, making bridging even more unlikely at large inter-surface separations.

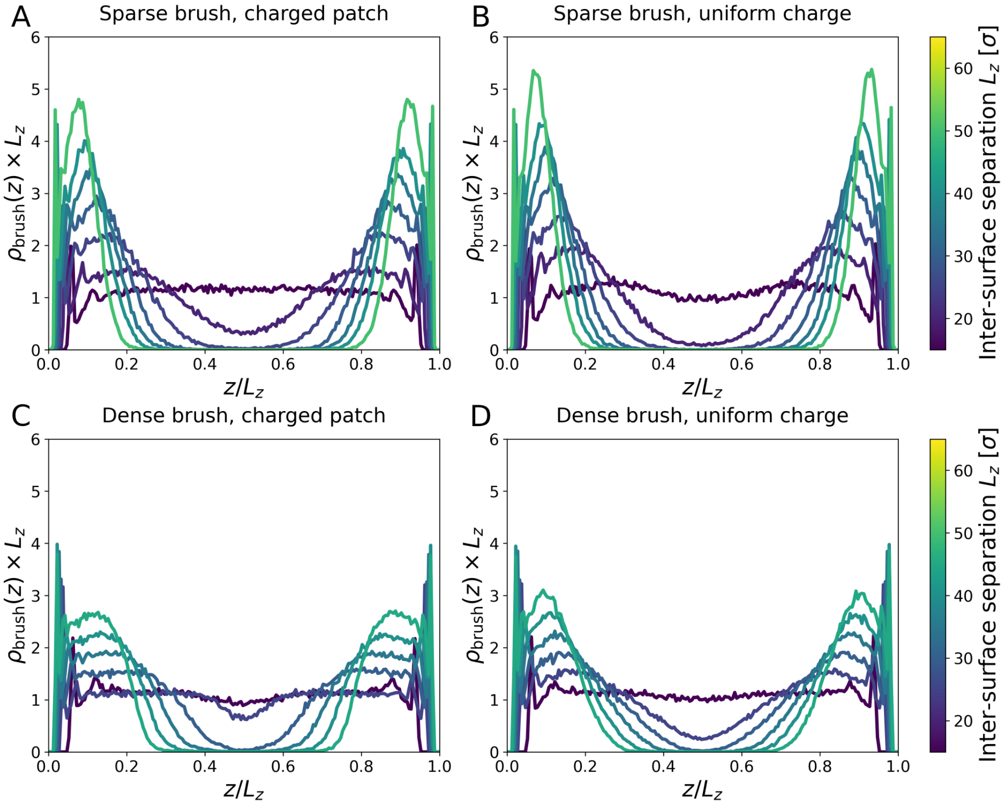


FIG. S12. Density profile of Ki-67 (brush polymer) along the $z$ axis. The horizontal axis was normalized by the inter-surface separation $L_z$. **(A)** Sparse brush, CP. **(B)** Sparse brush, uniform charge. **(C)** Dense brush, CP. **(D)** Dense brush, uniform.

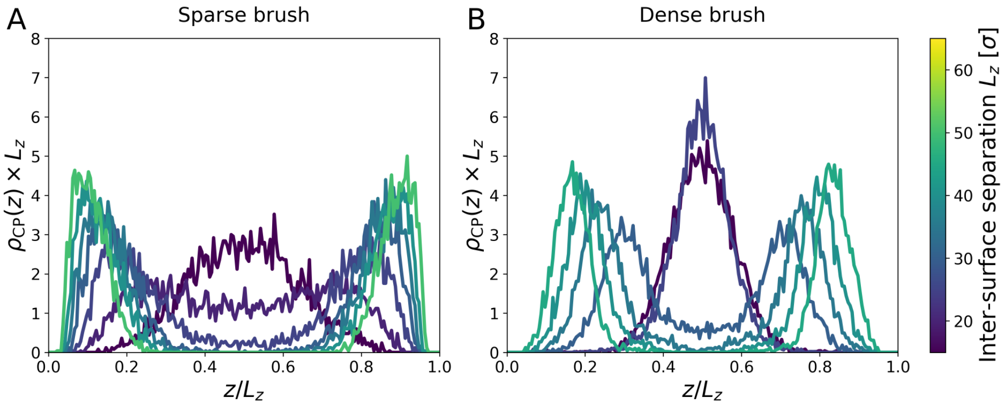


FIG. S13. Density profile of the CPs along the $z$ axis. The horizontal axis was normalized by the inter-surface separation $L_z$. **(A)** Sparse brush **(B)** Dense brush.

#### J. PMF dependence on the electrostatic interaction strength and screening length

In this Section, we test the robustness of our results when changing the parameters of the electrostatic interaction, Eq. S2. In particular, we vary the strength $K$ of the electrostatic interaction and the screening length $\lambda$. A decrease of $K$ would follow from an increase in the dielectric constant of the solvent, whereas a decrease of $\lambda$ would be caused by an increase in the ionic strength $I$ of the solvent.

In our simulations, the value of the parameter $K$ in Eq. (S2), controlling the strength of the electrostatic interaction, is $0.2\epsilon\sigma\mathcal{Q}^{-2}$. As our potential is only a coarse-grained, qualitative representation of electrostatic interactions, this is effectively a free parameter. In this Section, we show that the results presented in the

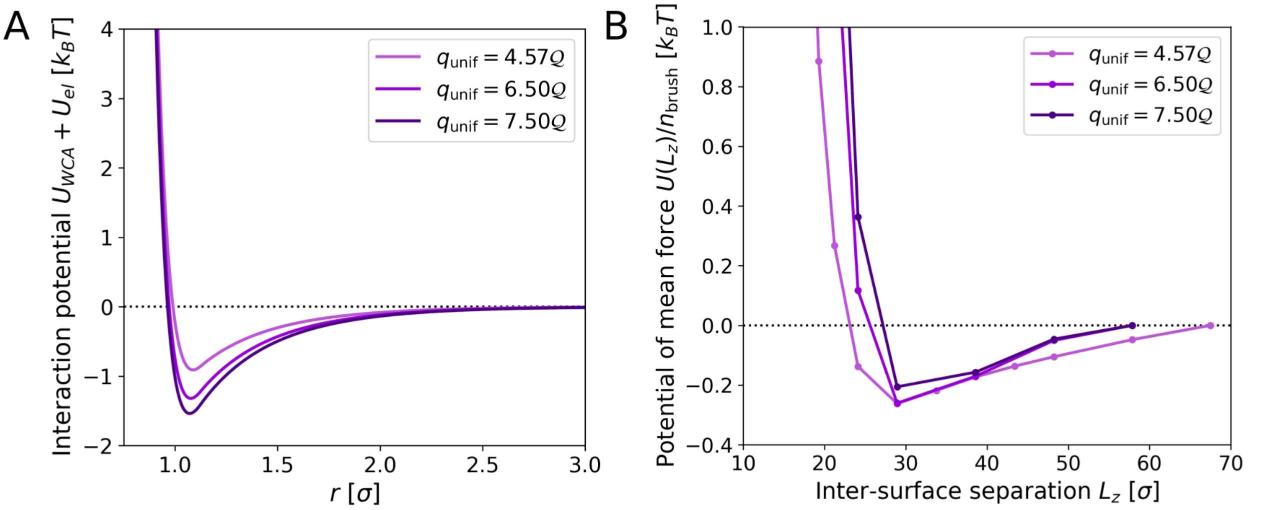


FIG. S14. (A) Total interaction potential $U_{\rm WCA} + U_{\rm el}$ [Eq. (S1)–(S2)] between RNA and uniformly charged brush polymers beads. (B) Potential of mean force (PMF) between the grafted surfaces for dense, uniformly charged brush with different values of the brush bead charge $q_{\rm unif}$. The value used throughout the rest of this work is $q_{\rm unif} = 4.57\mathcal{Q}$.

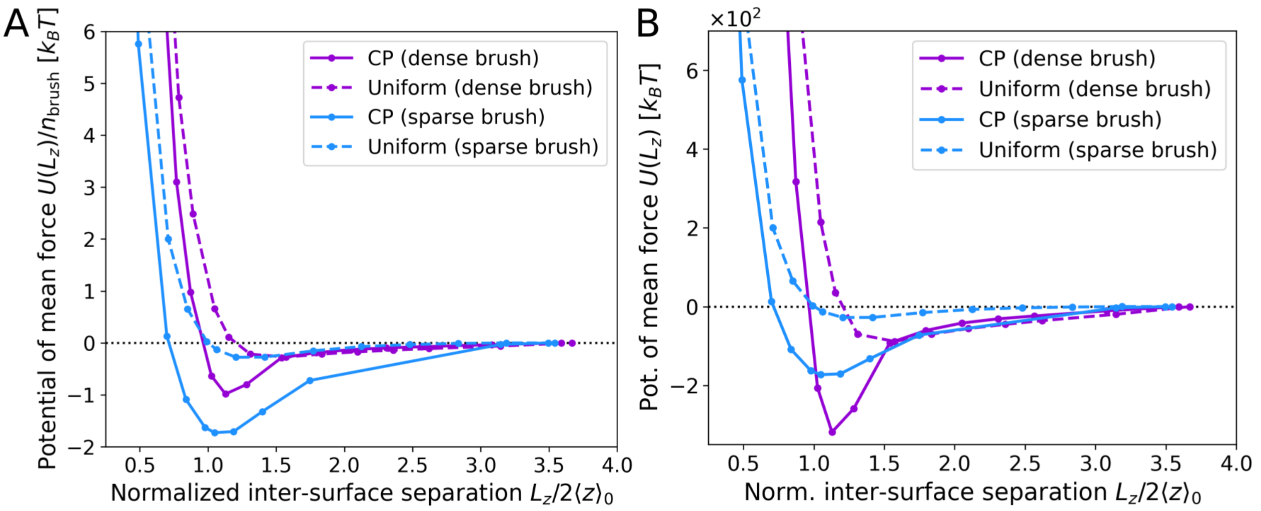


FIG. S15. Potential of mean force (PMF) between the grafted surfaces for strength of the electrostatic interaction $K = 0.1\epsilon\sigma\mathcal{Q}^{-2}$ [see Eq. (S2)]. This value is lower than the one used throughout the rest of this work, $K = 0.2\epsilon\sigma\mathcal{Q}^{-2}$, and corresponds to considering a solvent with a higher dielectric constant. **(A)**: PMF normalized by the number of polymers per brush $n_{\rm brush}$. **(B)**: Non-normalized PMF.

main text for the PMF remain valid when this parameter is decreased, which qualitatively corresponds to an increase of the dielectric constant of the solvent.

In Fig. S15A, we show the PMF, normalized by the number of Ki-67 polymers per brush $n_{\rm brush}$, for $K = 0.1\epsilon\sigma\mathcal{Q}^{-2}$. The range and depth of the attractive well of the normalized PMF change in a non-trivial way with the grafting density of the brush, analyzing which is out of the scope of the present work. We observe, however, that decreasing $K$ significantly reduces the electrostatic repulsion between the brushes. This has a larger impact on the sparse brush system, since for this one the excluded volume repulsion between the brushes is smaller. As a consequence, the attraction per brush polymer becomes stronger for the sparse brush than for the dense one. The total attraction, however, remains stronger for the dense brush, as shown in Fig. S15B, where we show the non-normalized PMF. However, our main result remains valid, in that the range and strength of the inter-surface attraction are always larger with the CP than with the

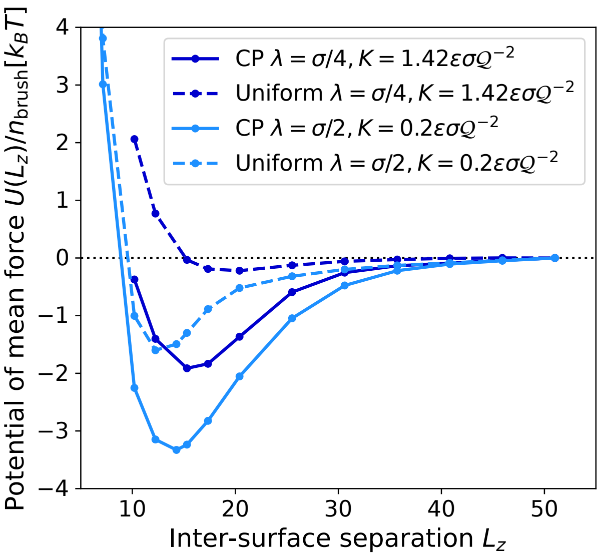


FIG. S16. Potential of mean force (PMF) between the grafted surfaces for the sparse brush. Here the potential used in the main text, with screening length $\lambda = \sigma/2$ and strength of the electrostatic potential $K = 0.2\epsilon\sigma\mathcal{Q}^{-2}$ (see Eq. S2), is compared to one with $\lambda = \sigma/4$ and $K = 1.42\epsilon\sigma\mathcal{Q}^{-2}$. With this choice of parameters, the electrostatic interaction for $r = \sigma$ is approximately the same for both potentials. The behavior of the PMF remains qualitatively the same, namely the CP leads to a significantly stronger attraction.

uniformly charged brush.

In Fig. S16, we compare for the sparse brush the potentials of mean force (PMF) for screening length $\lambda = \sigma/2$ (the value used in the main text, in light blue) and $\lambda = \sigma/4$ (dark blue). Here, we set $K = 1.42\epsilon\sigma\mathcal{Q}^{-2}$, such that the electrostatic interaction at contact, *i.e.* for $r = \sigma$, is unchanged. As one can see, we recover the same qualitative behavior observed in the main text: the attraction is significantly stronger for the CP case than for the uniformly charged case. The quantitative difference is likely due to the fact that fewer RNA beads interact on average with a brush polymer bead for the smaller screening length.

#### K. Subdiffusive dynamics of RNA monomers

In Fig. S17 we show the mean-squared displacement (MSD) of the RNA monomers for different systems. This quantity is defined as $r^2_{\rm m}(t) \equiv \langle|\mathbf{r}_i(t) - \mathbf{r}_i(0)|^2\rangle$, where $\mathbf{r}_i(t)$ is the position vector (at time $t$) of the $i$-th monomer. The data are compared to the monomer MSD for a freely diffusing RNA molecule (grey line), which is diffusive ($r^2_{\rm m}(t) \propto t$) approximately over the entire time range here considered. We recall that for a freely diffusing polymer in the absence of hydrodynamic interactions (as it is the case here), the Rouse model [13] predicts

$$\langle r^2_{\rm m}(t)\rangle \propto \begin{cases} t^{1/2} & t \lesssim \tau_R \\ t & t \gtrsim \tau_R, \end{cases} \tag{S30}$$

where $\tau_R \approx \zeta\sigma^2 N^2_{\rm RNA}/(k_BT)$ is the Rouse relaxation time, with $\zeta = 10m/\tau$ the viscous friction, $\sigma$ the monomer size and $N_{\rm RNA} = 100$ the number of beads in an RNA molecule. Thus, in our case, $\tau_R = 10^5\tau$. In the time range shown in Fig. S17, the monomer MSD for the free RNA molecule displays a broad crossover from a power law with slope 1/2 to one with slope 1.

The monomer dynamics in the presence of the brushes is markedly different: in the uniform charge case (B,D), we observe a marked Rouse-like subdiffusive regime with slope 1/2 that persists until very long times. In the main text, we have observed that the RNA diffuses as if the effective viscous coefficient was $\zeta_{\rm eff} = 100m/\tau$ instead of $\zeta = 10m/\tau$, due to being adsorbed onto the brush. We thus attribute this subdiffusive regime to a Rouse-like dynamics with a much larger relaxation time $\tau_R \approx \zeta_{\rm eff}\sigma^2 N^2_{\rm RNA}/(k_BT) = 10^6\tau$, which displaces to later times the crossover to diffusive dynamics.

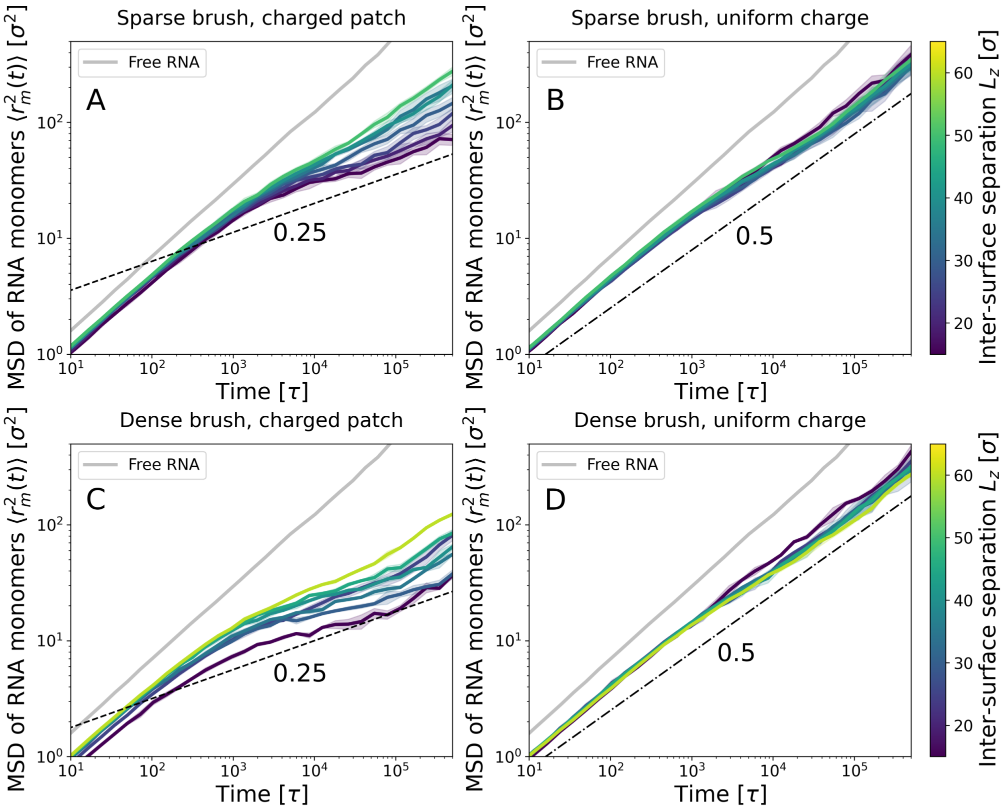


FIG. S17. MSD of RNA monomers for different systems. **(A)** Sparse brush, CP. **(B)** Sparse brush, uniform charge. **(C)** Dense brush, CP. **(D)** Dense brush, uniform. Grey curves: MSD of RNA monomers for freely diffusing RNA. Dashed lines: power law with slope 0.25. Dash-dotted line: power law with slope 1/2 (Rouse-like) [13].

The most interesting case, however, is that of the CP (A,C): here, instead of a subdiffusive regime with slope 1/2, we observe one with slope $\approx 0.25$. We attribute this much slower diffusion to the strong bonds between the RNA and the CPs: in order to diffuse, an RNA molecule has to break the bonds with many CPs, leading to a slow activated dynamics (sometimes called "sticky Rouse" dynamics) [14–22]. The exponent we find ($\approx 0.25$) is numerically close to the one observed for entangled polymers [5, 13], despite our system being unentangled (as proven by the fact that the dynamics of the center of mass of the RNA molecules is diffusive for uniformly charged brush). We note that, for unentangled melts of associative polymers, the exact value of the subdiffusive exponent has been shown to depend on the number of sticky spots on the chain and on the strength of the interaction [17].

#### L. Comparison to capillary forces

In this Section, we compare the scaling of the attractive force we measure with predictions for capillary forces. We highlight that, despite the qualitative similarities between the two types of interaction, we obtain a scaling of the pressure with distance which does not match the prediction for capillary forces. We thus conclude that our bridging mechanism is distinct from what expected for capillary interactions.

The capillary pressure between two infinite parallel plates can be computed from the Young-Laplace equation [23]:

$$P_{zz} = \frac{F_z}{A} \approx -\frac{\gamma}{R} = -\frac{2\gamma\cos(\theta)}{L_z}. \tag{S31}$$

where $\gamma$ is the surface tension, $R$ is the radius of curvature of the meniscus and $\theta$ is the contact angle [24, 25]. In Fig. S18A, we show the $zz$ component of the pressure tensor for the CP case. We attempted to fit the data with the function

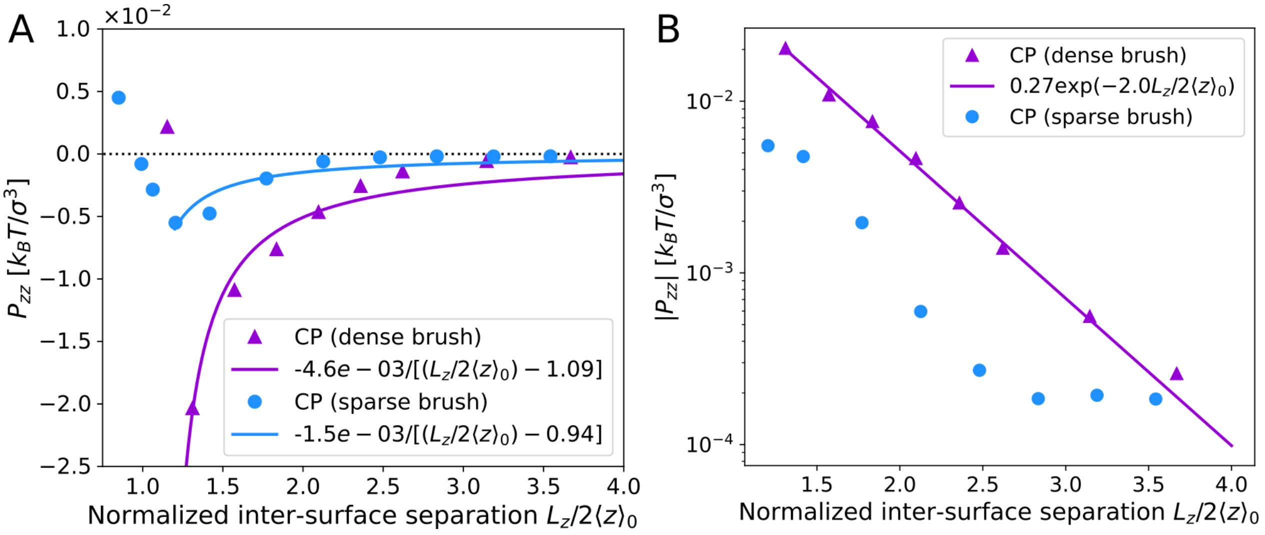


FIG. S18. **(A)** Vertical ($zz$) component of the pressure tensor (pressure between the plates) as a function of the normalized inter-surface separation for the brushes with CP. Symbols: simulation data for the dense (triangles) and sparse (circles) brush. Solid lines: best fit of the data with Eq. (S32). **(B)** Absolute value of the pressure between the plates as a function of the normalized inter-surface separation. Only data for the increasing part of $P_{zz}(L_z)$ are shown. Symbols as in (A). Solid line: exponential fit of the data for the dense brush.

$$P_{zz}(L_z) = \frac{A}{(L_z/2\langle z\rangle_0) - B}, \tag{S32}$$

where $A$ has units $k_BT\sigma^{-3}$ and the factor $B$ is to take into account that in our case the two brushes come into contact for $L_z/2\langle z\rangle_0 \approx 1$ (thus, we expect $B \approx 1$, which is confirmed by the fit results). The solid lines represents the results of the fit: One can see that the agreement with Eq. (S32) is rather poor. Attempting to fit the data for the uniformly charged brush gives an even poorer results (not shown). Thus, we conclude that the scaling of the force with distance differs from what predicted for capillary interactions.

In Fig. S18B, we show that the increasing part of $P_{zz}$ is much better fitted by an exponential. To show this, we plot the absolute value of the pressure, $|P_{zz}|$, in range of inter-surface separations for which $P_{zz}$ increases. One can see that, for the dense brush, $|P_{zz}|$ is well fitted by an exponential:

$$|P_{zz}| = 0.27k_BT\sigma^{-3}\exp(-2.0L_z/2\langle z\rangle_0) \approx 0.27k_BT\sigma^{-3}\exp(-L_z/\langle z\rangle_0) \tag{S33}$$

The data for the sparse brush also show an approximately exponential decrease, but the agreement is not as good at large inter-surface separations. The observation that the decay is approximately exponential matches qualitatively with our theoretical prediction that the number of bridges decreases exponentially with inter-surface separation, see Eq. (2) and Fig. 4 in the main text, and that it is bridging that drives attraction.

Our conclusion is thus that, despite the similarities between the attraction mechanism we describe and capillary forces, the physical details are not the same as those predicted for continuum fluids. We attribute this to the fact that the scales of the molecules leading to the binding mechanism (RNA *vs* water) are very different, and to the fact that RNA-mediated attraction has an important entropic component. Finally, we mention that, in the cellular context, additional layers of complexity may exist – for example, soluble Ki-67 or other factors could contribute to inter-chromosomal interactions or influence their spatial organization through mechanisms not accounted for in the model. In this context, capillary interactions might play a relevant role.

---

[10] M. Rubinstein and R. H. Colby, *Polymer physics* (Oxford University Press New York, 2003).

[23] P.-G. De Gennes, F. Brochard-Wyart, and D. Quéré, *Capillarity and wetting phenomena: drops, bubbles, pearls, waves* (Springer Science & Business Media, 2003).

[24] B. Gouveia, Y. Kim, J. W. Shaevitz, S. Petry, H. A. Stone, and C. P. Brangwynne, Capillary forces generated by biomolecular condensates, Nature **609**, 255 (2022).

[25] S. Cheng and M. O. Robbins, Nanocapillary adhesion between parallel plates, Langmuir **32**, 7788 (2016).
